# Transient proliferation by reversible YAP and mitogen-control of the cyclin D1/p27 ratio

**DOI:** 10.1101/2024.10.11.617852

**Authors:** Katherine R. Ferrick, Yilin Fan, Nalin Ratnayeke, Mary N. Teruel, Tobias Meyer

**Affiliations:** Department of Cell and Developmental Biology, Weill Cornell Medicine, New York, NY, USA; Department of Chemical and Systems Biology, Stanford Medicine, Stanford, CA, USA; Department of Pathology and Center for Cancer Research, Massachusetts General Hospital and Harvard Medical School, Boston, MA, USA; Cancer Biology and Genetics Program, Sloan Kettering Institute, Memorial Sloan Kettering Cancer Center, New York, NY, USA; Department of Biochemistry, Weill Cornell Medicine, New York, NY, USA

## Abstract

Hippo-YAP signaling orchestrates epithelial tissue repair and is therefore an attractive target in regenerative medicine. Yet it is unresolved how YAP integrates with mitogen signaling and contact inhibition to control the underlying transient proliferative response. Here we show that reduced contact inhibition, increased mitogen signaling, and YAP-TEAD activation converge on increasing the nuclear cyclin D1/p27 protein ratio during G1 phase, towards a threshold ratio that dictates whether individual cells enter or exit the cell cycle. YAP increases this ratio indirectly, in concert with mitogen signaling, by increasing EGFR and other receptors that signal primarily through ERK. After a delay, contact inhibition suppresses YAP activity which gradually downregulates mitogen signaling and the cyclin D1/p27 ratio. Increasing YAP activity by ablating the suppressor Merlin/NF2 reveals a balancing mechanism in which YAP suppression and contact inhibition of proliferation can be recovered but only at higher local cell density. Thus, critical for tissue repair, robust proliferation responses result from the YAP-induced and receptor-mediated prolonged increase in the cyclin D1/p27 ratio, which is only reversed by delayed suppression of receptor signaling after contact inhibition of YAP.

## INTRODUCTION

Acute chemical and physical injuries inactivate the Hippo pathway, resulting in transient activation of the transcriptional cofactor YAP to increase proliferation, induce metabolic changes, and promote matrix remodeling^1,2^. YAP activation and nuclear translocation initiates transcription through binding to TEAD and other nuclear transcription factors^3^. Though YAP and its paralog TAZ are dispensable for the basal proliferation of several tissues, they are crucial for regeneration and repair following injury in tissues such as skin, liver, and kidney^1^. Inhibitors of Hippo pathway proteins are therefore being developed for potential clinical use in tissue repair^4–6^. While the core genes regulating the Hippo pathway are well-established, the complexity of the downstream signaling processes has made it difficult to resolve the molecular mechanism of the transient proliferation response using animal models.

Hippo signaling is kept active in epithelial sheets and other tissues by cell-cell contacts, such as adherens and tight junctions, and is inactivated by mechanical stress during tissue repair^1,7,8^. Adaptor proteins like Merlin (NF2) contribute to contact inhibition signaling to activate LATS1/2, which is the principal kinase of the Hippo pathway suppressing YAP-TEAD activity through multisite phosphorylation^9,10^. Consequently, loss of Merlin or LATS activity leads to dramatic increases in the size of tissues and organs, and tumorigenesis^9,11,12^. Despite the general role of YAP activation in promoting proliferation, the mechanism by which YAP integrates with other signaling pathways to regulate both cell cycle entry and exit is unresolved.

Different mechanisms have been described through which YAP signaling could increase proliferation. Constitutive YAP activation increases expression of a broad array of cell cycle genes, including DNA replication factors and regulators of G2/M progression^13–15^. YAP also promotes secretion of various signaling regulators^16,17^. However, cell cycle entry is not only controlled by YAP, but is also regulated by both receptor signaling and contact inhibition^18,19^. Ultimately, cell cycle entry is controlled by cyclin-dependent kinase (CDK) activation in G1 phase which must phosphorylate and inactivate retinoblastoma protein (Rb), the suppressor of the cell cycle transcription factor E2F. As persistent YAP activation induces organ hyperplasia and cancer, YAP-TEAD must be robustly inactivated after its initial induction, once the tissue repair is completed. Thus, to understand tissue repair, it is crucial to determine how YAP, mitogens, and contact inhibition control Rb phosphorylation to first promote and then suppress cell cycle entry.

Here we use multiplexed, single-cell immunofluorescence analysis to examine YAP-regulated proliferation in thousands of epithelial cells in a monolayer. We show that over a wide range of stimuli, proliferation is controlled through a balance mechanism whereby mitogen signaling is opposed by contact inhibition of YAP. Strikingly, mitogen signaling, contact inhibition, and YAP-activation converge on the cyclin D1/p27 ratio during G1 phase, which dictates whether Rb is phosphorylated and cells enter the cell cycle. We show that YAP activation increases the levels of EGFR and other surface receptors, which in turn signal primarily through MEK-ERK to increase the cyclin D1/p27 ratio. We further identify the mechanism that prolongs the proliferation response: acute inactivation of YAP gradually reduces EGFR signaling to lower the cyclin D1/p27 ratio but only after a delay. Moreover, we show that YAP signaling can be robustly terminated: even a strong increase in YAP activity generated by ablation of the upstream Hippo component Merlin could still be reversed by contact inhibition of YAP, but only at higher local cell density. Together, our study identifies a YAP-controlled balancing mechanism between contact inhibition and receptor signaling that drives an increase and delayed decrease in the cyclin D1/p27 ratio. The delay in the balance ensures that the YAP-induced proliferation response is prolonged to allow for sufficient cell divisions while the robust YAP inactivation ensures that the proliferation response is inherently transient.

## RESULTS

### Contact inhibition of YAP opposes mitogen stimulation in a balance mechanism controlling cell proliferation

To assess mitogen and density-dependent changes in proliferation, we measured both phosphorylation of retinoblastoma protein (p-Rb, S807/811), which is a bimodal marker of cyclin-CDK activity^20^ that is low in arrested cells and remains high in cycling cells from G1 through mitosis, and EdU incorporation into DNA, which occurs during DNA replication in S phase (Figure 1A, S1A). We determined how YAP activity changes the mitogen dependence of proliferation by stably expressing a doxycycline (DOX)-inducible, constitutively active YAP variant (5 regulatory serines mutated to alanine^21^, CA-YAP) in hTERT-immortalized human retinal pigmented epithelial cells (RPE1), which reversibly arrest the cell cycle with serum starvation or contact inhibition^22^. As leaky expression of CA-YAP (without DOX induction) was sufficient to increase proliferation (at a similar expression level compared to the endogenous YAP, Figure S1B), we did not use DOX induction unless otherwise stated.

**Figure 1.**
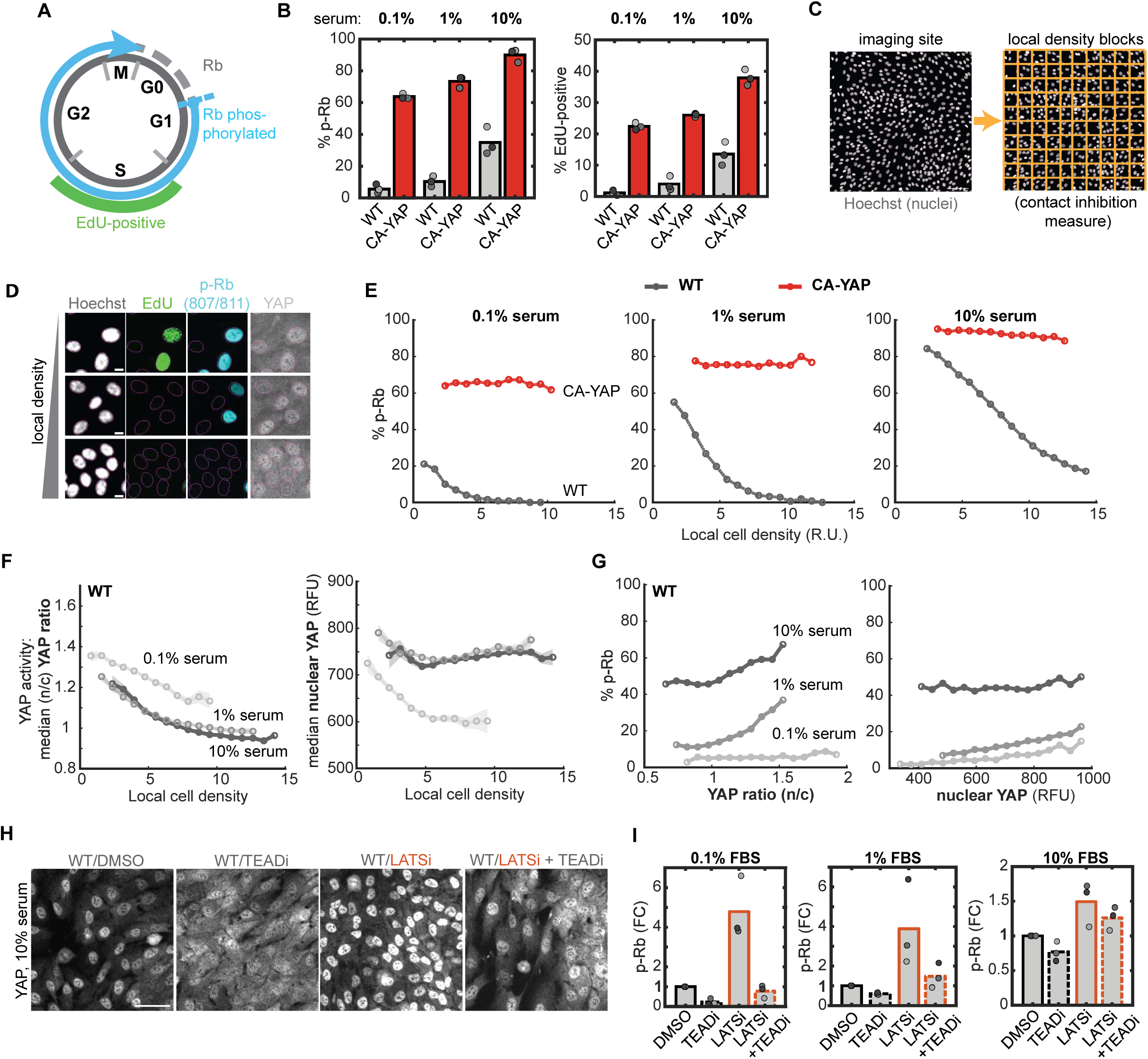
Contact inhibition of YAP opposes mitogen stimulation in a balance mechanism that controls proliferation. **A)** Diagram of cell cycle phases: EdU incorporation marks S-phase, Rb hyperphosphorylation (p-Rb, S807/811) cycling cells. **B)** Mean percent p-Rb (left) and EdU-positive (right) for wild-type (WT) and CA-YAP cells maintained in increasing serum levels (0.1%, 1%, or 10%) for 36h. N = 3 independent experiments. Student’s t-test (p-Rb, EdU): p= 6.86×10^-6^, 1.01×10^-5^ (0.1% serum), p= 2.50×10^-5^, 8.13×10^-5^ (1% serum), p= 5.90×10^-4^, 7.13×10^-4^ (10% serum). **C)** Representation of local cell density metric, by subdividing imaging sites into 10×10 grids and counting the cells (by centroid of segmented nucleus) for each block (see also Methods, scale bar = 100 μm). **D)** Sample images of RPE1 cells at increasing local cell density: Hoechst for DNA (white), EdU for S-phase cells (green), p-Rb for cycling cells (blue), and YAP (grey). Magenta outline is the automated nuclear mask (scale bar = 10 μm). **E)** Percent p-Rb binned by local density for WT and CA-YAP treated as in (B). The percent was calculated as the fraction of p-Rb-positive cells from cells binned by the local cell density metric (relative units, R.U.), n > 20,000 cells/condition. See also **Figure S1**. **F)** Median nuclear/cytoplasmic (n/c) YAP ratio (left) and nuclear YAP levels (right, real fluorescent units, RFU) per local density bin for WT cells treated as in (C), n > 20,000 cells/condition. Shaded error bars are 95% confidence intervals. **G)** Percent p-Rb for WT cells binned by their YAP ratio (left) or nuclear YAP levels (right) for cells treated as in (C). **H)** Representative images of YAP staining for WT cells treated with TEAD inhibitor (TEADi, 0.5 μM GNE-7883), LATS inhibitor (LATSi, 0.5 μM TDI-011536) or in combination in 10% serum for 24h. **I)** Fold-change in percent p-Rb for WT cells treated with TEADi, LATSi, or in combination, normalized to DMSO in 0.1% (left), 1% (middle), or 10% serum (right). N = 3 independent experiments. All binned plots have n>100 cells/bin and >20,000 cells/condition.

Using increasing concentrations of serum (fetal bovine), we found that mean levels of proliferation (by both p-Rb and EdU, Figure 1B) were consistently higher in CA-YAP relative to wild type (WT) cells at all mitogen levels and serum-independent, as previously reported^23^. However, even within epithelial monolayers plated at identical densities and subject to the same mitogen stimuli, there is considerable heterogeneity in cell cycle signaling^22,24,25^. We therefore used single-cell analysis to distinguish between various plausible mechanisms of how YAP activation might be linked to both contact inhibition and mitogen signaling to control cell cycle entry. We use the terms “contact inhibition” and “local cell density” interchangeably since the local cell density in monolayers closely predicts whether proliferation is suppressed in individual RPE1 cells^22^. We quantified local cell density by subdividing segmented nuclei for each image into a 10 x 10 grid (∼75 x 75 μm) and counting the cells within each block (Figure 1C). Each cell was then assigned its local density based on the number of cells in its block (Figure S1C, see also Methods) and compared with the fraction of proliferating cells (Figure 1D).

In WT cells, the proportion of proliferating cells progressively decreased as local cell density increased for high and low serum levels, supporting that mitogen signaling and local contact inhibition oppose each other in regulating proliferation (Figure 1E, S1D). This conclusion was also supported by an alternative metric of local cell density based on the area of neighboring cell nuclei within a radius of similar scaling^22^ (Figure S1E).

Strikingly, even with weak YAP expression, the cell density dependence of proliferation was abolished in CA-YAP cells regardless of mitogen levels (Figure 1E), demonstrating that contact inhibition of proliferation requires contact inhibition of YAP activity. Moreover, this confirmed that CA-YAP cells retained mitogen sensitivity independent of contact inhibition since increasing serum levels further increased the percentage of proliferating cells. Thus, the inverse regulation between mitogens and cell density represents an intuitive balancing mechanism: the inhibition of proliferation at a given cell density can be overcome by increasing mitogen signaling, and the increase in proliferation after increasing mitogen signaling can be antagonized once cell division has increased local cell density. However, persistent activation of YAP breaks the balance and makes proliferation independent of cell density.

We next tested whether contact inhibition and mitogen signaling regulate endogenous YAP activity. As the ratio of the nuclear to cytoplasmic (n/c) YAP and the absolute nuclear level of YAP are both relative measures of YAP activity^26,27^, we measured both parameters as a function of local cell density (Figure 1F). At 0.1% serum, we observed the expected inhibition of the YAP n/c ratio and nuclear levels at increasing local density. At 1% and 10% serum, the YAP n/c ratio decreased at higher cell densities, but the nuclear level of YAP was higher than for low serum conditions. We used this single-cell data to analyze the percentage of cells with phosphorylated Rb as a function of the relative YAP activity. Unexpectedly, using both metrics, increasing serum levels increased proliferation for cells with identical YAP levels (Figure 1G). Thus, while YAP activation and mitogen stimuli synergistically increase proliferation, we find that mitogens can also increase proliferation directly without increasing YAP levels or localization.

Finally, we determined how acute increases and decreases in YAP activity regulate Rb phosphorylation. Using recently developed inhibitors, we inhibited YAP-TEAD signaling with a pan-TEAD inhibitor^28^, and activated endogenous YAP using a LATS1/2 inhibitor^5,29^. We confirmed that TEAD inhibition promoted a strong cytoplasmic localization of endogenous YAP while LATS inhibition triggered its nuclear enrichment (Figure 1H). TEAD inhibition greatly reduced the fraction of proliferating cells for low but not for high serum stimuli, suggesting that YAP-TEAD activity is particularly critical for inducing proliferation for low mitogen stimulation conditions (Figure 1I, S1F). Conversely, LATS inhibition greatly boosted proliferation at low serum stimuli and had a smaller relative effect at high serum stimuli due to the already high percentage of proliferating cells (Figure S1G). Finally, this analysis revealed that high mitogens can induce proliferation independently of YAP-TEAD activity since TEAD inhibition could reverse the portion of proliferation induced by YAP but not a YAP-independent component of mitogen-induced proliferation (Figure 1I, Figure S1H).

We conclude that contact inhibition of YAP activity and mitogen signaling compete to respectively suppress or promote proliferation over a wide range of cell densities and mitogen stimuli. However, this balance can be broken in two ways: by high mitogen stimuli inducing proliferation in cells lacking YAP-TEAD activity and persistent activation of YAP abolishing contact inhibition of proliferation.

### YAP-regulated cell cycle entry and exit is dictated by the cyclin D1/p27 threshold

We next used the quantitative analysis of the balance between contact inhibition and mitogen signaling to identify the mechanism of how YAP regulation controls cell-cycle entry and exit. Cells enter the cell cycle during the G1 phase by activating cyclin-CDK complexes to hyperphosphorylate Rb and derepress the E2F cell cycle transcription program^20^. In contrast, in cells that exit the cell cycle or stay arrested, cyclin-CDKs stay inactive and Rb remains dephosphorylated. In monolayers of RPE1 and other epithelial cells, this bimodal Rb phosphorylation is thought to be primarily controlled by the CDK4/6 activator cyclin D1 and the cyclin-CDK2/4/6 inhibitor p27^22^, and cyclin D1 is generally thought to contribute to the YAP-mediated proliferation regulation^3^. We included p27 in our analysis since both cyclin D1^30^ and p27 are key regulators of cell proliferation in vivo, with p27 deletion resulting in multi-organ hyperplasia^31^.

Using image-based cytometry and multiplexed immunofluorescence^32^, we first computationally gated for G1 cells by their diploid 2N DNA content and lack of incorporation of EdU (Figure S2A). We then measured the levels of nuclear localized cyclin D1 and p27 and whether Rb was phosphorylated in each G1-phase cell. Notably, p27 protein levels are often used as a marker of non-proliferating cells, but we observed substantial heterogeneity in the nuclear levels of both p27 and cyclin D1 across individual cells irrespective of Rb phosphorylation (Figure S2B). Increasing the mitogen stimulus (from 0.1% to 10% serum) resulted in the cell population shifting towards higher average cyclin D1 levels and lower average p27 levels.

We next determined how local cell density and YAP activity regulate the cyclin D1 and p27 levels. As local density increased, the median cyclin D1 level of WT cells gradually decreased, while the median p27 level strongly increased (Figure 2A). This trend was opposite to the effects of increasing serum, which both globally increased cyclin D1 levels and suppressed the density-dependent increase in p27 protein. Markedly, the CA-YAP cells lost the serum and density dependence of both, keeping the levels of cyclin D1 high and p27 low in G1 phase, arguing that YAP activity is not only upregulating the level of cyclin D1 but also downregulating the level of p27 (Figure 2B).

**Figure 2.**
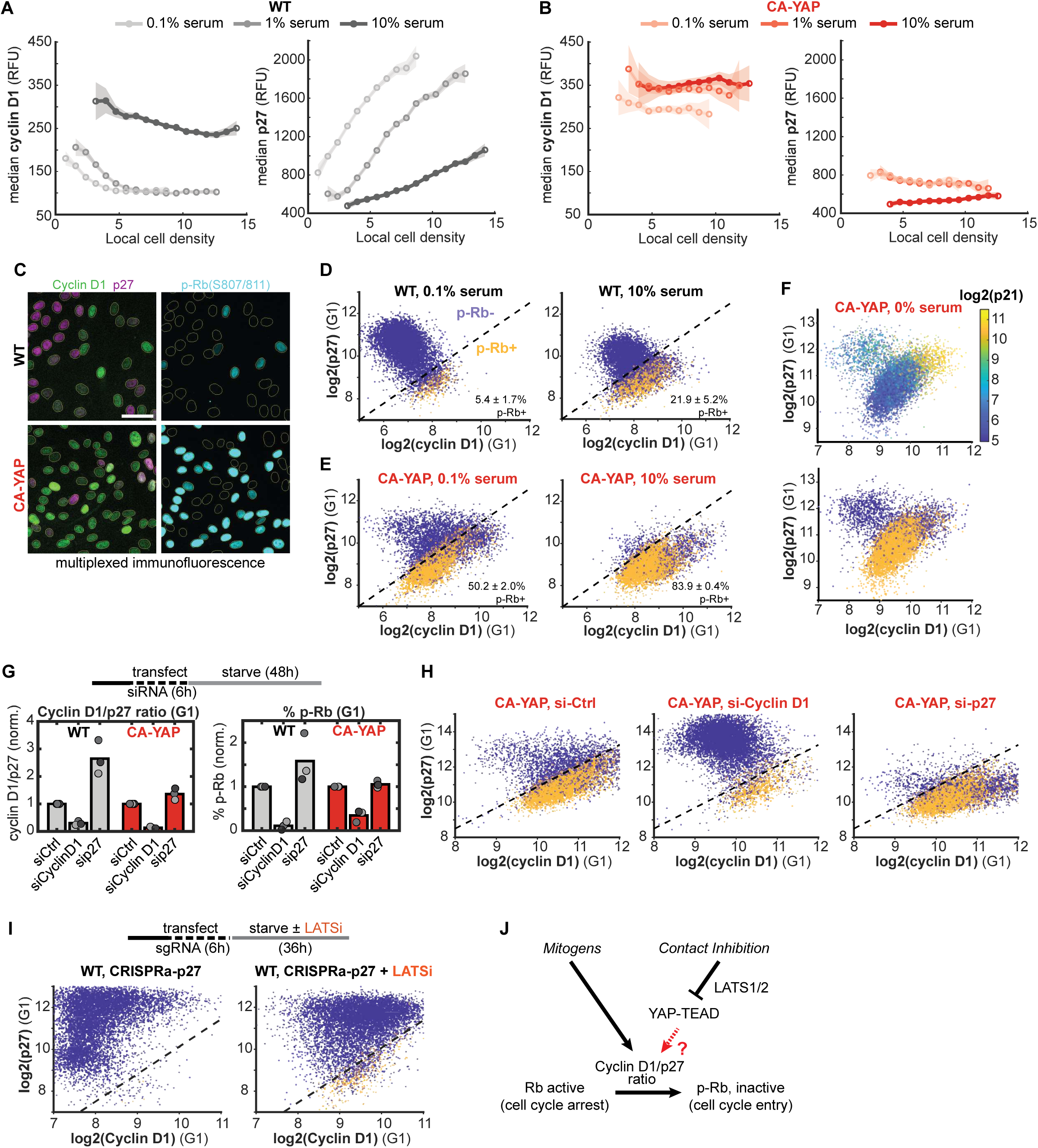
YAP-regulated cell cycle entry and exit is dictated by the cyclin D1/p27 threshold. **A-B)** Median levels (real fluorescent units (RFU)) of cyclin D1 (left) and p27 (right) per local cell density bin for WT (A) and CA-YAP (B) cells at 0.1%, 1%, and 10% serum. Binned data only include G1-gated cells, n > 100 cells/bin. Shaded error bars are 95% confidence intervals, n > 15,000 cells per condition. Data are representative of 2 independent experiments. **C**) Representative images of multiplexed immunofluorescence staining comparing cyclin D1 (green) and p27 levels (magenta) with aligned p-Rb stain (cyan, with gray nuclear mask overlay) in WT or CA-YAP cells. Scale bar = 50 μm. **D-E**) Representative single-cell scatterplots of co-stained cyclin D1 and p27 levels (log2-transformed), colored by p-Rb (purple, p-Rb-negative; orange, p-Rb-positive) in WT (D) and CA-YAP (E). Left, 0.1% serum; right, 10% serum. Mean ± SD of 2 independent experiments. **F)** Representative single-cell scatterplots of serum-starved CA-YAP cells co-stained for cyclin D1 and p27 protein levels, colored by p21 protein levels (<u>top:</u> parula color scale) or p-Rb status (<u>bottom</u>: purple, p-Rb-negative; orange, p-Rb-positive). **G)** <u>Top</u>: experimental conditions for siRNA transfection followed by 48h serum starvation. <u>Bottom</u>: mean cyclin D1/p27 ratio (left) and p-Rb (right) for WT and CA-YAP cells transfected with siRNA against cyclin D1 and p27, normalized to non-targeting control (siCtrl). N = 3 independent experiments. **H)** Representative single-cell scatterplots of co-stained cyclin D1 and p27 levels colored by p-Rb for CA-YAP cells treated as in (G). **I)** <u>Top</u>: experimental conditions for sgRNA-mediated overexpression (CRISPRa) of p27 in dCas9-expressing WT cells treated with DMSO or LATSi (0.5 μM TDI-011536). <u>Bottom</u>: Representative single-cell scatterplots of co-stained cyclin D1 and p27 levels colored by p-Rb for p27-overexpressing cells treated with DMSO (left) or LATSi (right). **J)** Schematic of mitogen signaling and contact inhibition of YAP-TEAD activity converging on the cyclin D1/p27 ratio to control whether cells enter or exit the cell cycle. Dashed lines approximate separation of p-Rb-positive and p-Rb-negative cell populations. Single-cell data are representative of 2 independent experiments, with n = 8,000 random cells/scatter plot and gated for G1-phase (defined as 2N DNA/EdU-negative status). See also **Figure S2**.

Explaining why the level of cyclin D1 or p27 alone cannot predict whether Rb is phosphorylated, only the level of cyclin D1 relative to p27 was tightly correlated with Rb phosphorylation in individual cells (Figure S2C). Only cells with relatively more cyclin D1 than p27 had Rb phosphorylated (Figure 2C-D). Thus, even for low mitogen levels, a small fraction of cells had sufficiently high cyclin D1 and low p27 to have Rb phosphorylated. Conversely, at high mitogen stimulation, more cells populated the high-cyclin D1/low-p27 state and had Rb phosphorylated, demonstrating a critical threshold cyclin D1/p27 ratio for proliferation.

Strikingly, in CA-YAP cells with persistently active YAP, Rb phosphorylation remained closely predicted by the relative nuclear cyclin D1 and p27 levels (Figure 2E, S2D-E), implying that YAP is signaling through both cyclin D1 and p27 to control proliferation. Notably, we identified some CA-YAP cells with high cyclin D1 that did not have Rb phosphorylated, but these cells had increased protein levels of another CDK inhibitor, p21, which we demonstrated in separate experiments (Figure 2F). Since p21 levels were only weakly regulated by serum starvation or contact inhibition (Figure S2F-G), p27 is functioning as the primary proliferation suppressor relevant for integrating mitogen signaling, contact inhibition, and YAP signaling^33,34^.

The fundamental role of the cyclin D1/p27 ratio in YAP-regulated proliferation is further seen in a comparison of perturbations that change the cyclin D1/p27 ratio and correspondingly change Rb phosphorylation (Figure 2G). The siRNA-mediated knockdown of cyclin D1 strongly reduced the proportion of cells with phosphorylated Rb by shifting most of the CA-YAP cells to a high-p27 and quiescent state, but again preserved the control of Rb phosphorylation by the relative cyclin D1/p27 ratio (Figure 2H, S2H). Moreover, knocking down p27 in WT cells resulted in more cells ending up below the dashed line but still preserved the control of Rb phosphorylation by the cyclin D1/p27 ratio (Figure 2G). Conversely, in cells where p27 levels were increased by CRISPRa induction, most cells were above the relative threshold line and proliferation was completely suppressed (Figure 2I). However, LATS inhibitor-mediated activation of YAP in these cells lowered p27 and increased cyclin D1 to push some cells below the threshold cyclin D1/p27 ratio where they could again proliferate (Figure 2I).

We conclude that the crucial parameter for the YAP-regulated control of cell cycle entry is the nuclear cyclin D1/p27 ratio in G1 phase, with a threshold dictating whether individual cells enter or exit the cell cycle (Figure 2J). Moreover, we show that there is a general convergence of contact inhibition, mitogen signaling, and YAP activity, since all act through the same downstream cyclin D1/p27 ratio. Notably, we were only able to identify this molecular mechanism of how YAP controls the cell cycle entry and exit decision through multiplexed measurements of p27, cyclin D1 and p-Rb levels within the same, G1-gated single cell.

### YAP regulation gradually changes the nuclear cyclin D1/p27 ratio

We next used RNA sequencing (RNAseq) to distinguish between different potential signaling pathways through which YAP activation may reversibly regulate cyclin D1 and p27 levels over time. Previous studies suggested various regulatory mechanisms, such as direct induction of cyclin D1^14^, indirect induction of mitogen signaling^16,35^, or reprogramming the cell state^15,36,37^. We measured gene expression changes up to 36-hours following CA-YAP induction in low serum cells (Figure 3A). Critically, we inhibited CDK4/6 activity in these cells (palbociclib) to keep cells in a state with minimal E2F activation to suppress the otherwise dominating effects of the E2F cell cycle transcription program that may obscure the YAP-specific transcription program.

**Figure 3.**
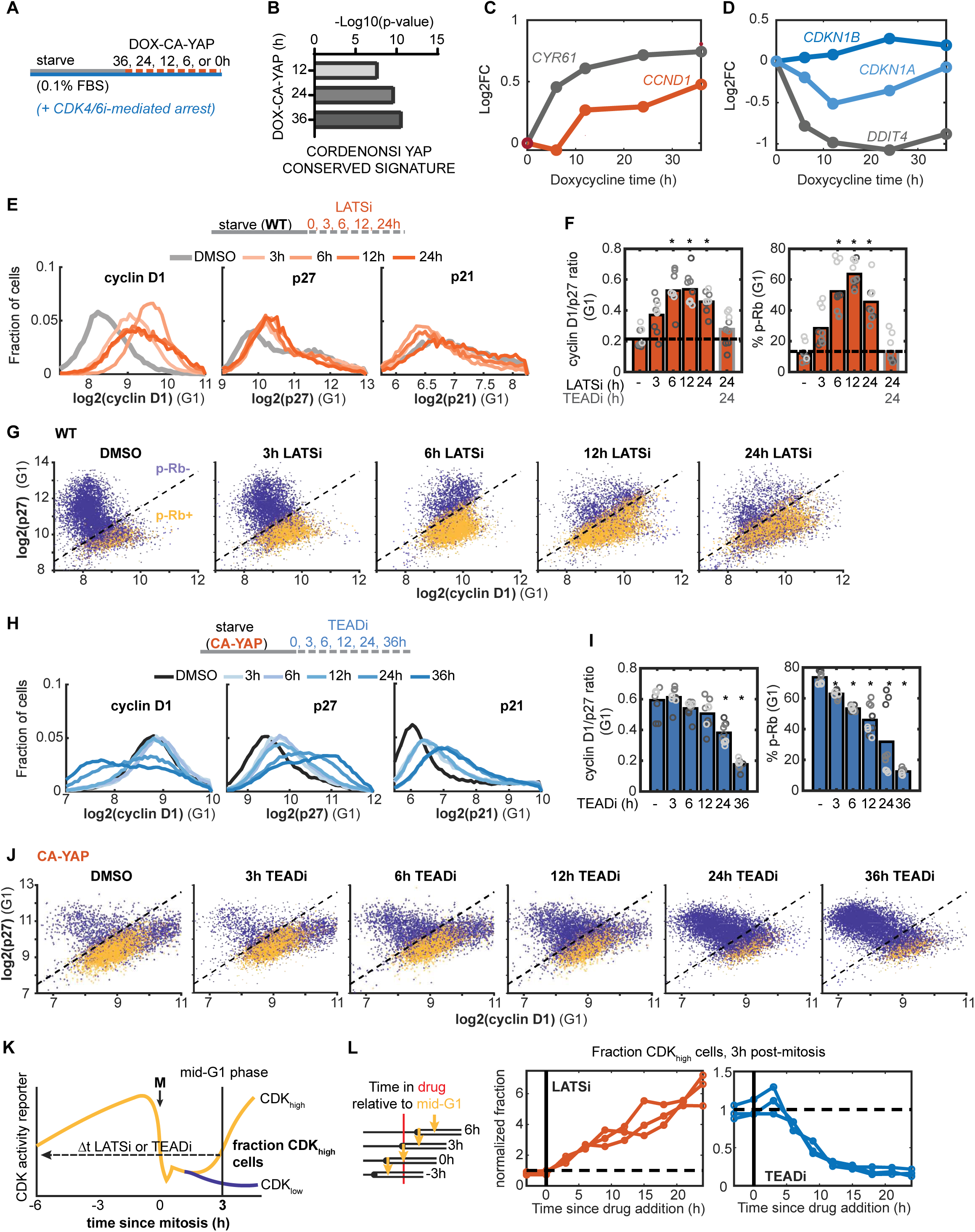
Delayed YAP-mediated control of the cyclin D1/p27 ratio and cell cycle arrest. **A)** Schematic for CA-YAP expression in RPE1 cells. RNA-sequencing performed in confluent cells pre-conditioned at 0.1% and palbociclib-arrested (CDK4/6i, 1 μM), prior to doxycycline-induction over 36h (0.5 μg/mL DOX). See also **Figure S3** and Methods. **B)** Up-regulated genes (q<0.05, Log2 fold-change >1) for 12, 24, and 36h YAP induction showed the Cordenonsi YAP signature (using ToppGene)^38^. **C-D)** Log2 fold-change (Log2FC) induction (C) of *CYR61* (canonical YAP target gene, grey) compared with *CCND1* (cyclin D1, red) and (D) regulation of DDIT4 (suppressed YAP target gene, grey), *CDKN1A* (p21, dark blue), and *CDKN1B* (p27, blue) at times since overexpression. Adjusted p-values (NS at all other time points): *CCND1*, 3.17×10^-4^ (12h), 4.45 x10^-5^ (24h), 3.96 x10^-12^ (36h); *CDKN1A*, 8.65×10^-7^ (12h), 9.5×10^-4^ (24h); *CDKN1B*, 0.0012 (24h), 0.0312 (36h). **E)** <u>Top</u>: Experimental conditions for LATSi treatment (0.5 μM TDI-011536) of serum-starved WT cells for 0-24h. <u>Bottom</u>: Histogram distribution of cyclin D1, p27, and p21 protein levels over LATSi time course. **F)** Mean cyclin D1/p27 ratio (left) and percent p-Rb (right) for cells treated as in (E). N = 3 independent experiments and 3 well-replicates/experiment, as indicated by matched colors. Student’s t-test: cyclin D1/p27, p = 0.0062 (6h), 0.0022 (12h), 2.63×10^-4^ (24h); p-Rb, p = 0.0217 (6h), 4.12×10^-4^ (12h), 0.0087 (24h). **G)** Representative single-cell scatterplots of cyclin D1 and p27 protein levels colored by p-Rb status (purple, p-Rb-negative; orange, p-Rb-positive). **H)** <u>Top</u>: experimental conditions for TEADi treatment (1 μM GNE-7883) of serum-starved CA-YAP cells. <u>Bottom</u>: Histogram distribution of cyclin D1, p27, and p21 protein levels over TEADi time course. **I)** Mean cyclin D1/p27 ratio (left) and percent p-Rb (right) for cells treated as in (H). N = 3 biological replicates, where well-replicates are indicated by matched colors. Student’s t-test: cyclin D1/p27 ratio, p = 0.027 (24h), p = 0.0015 (36h); % p-Rb, p = 0.0188 (3h), p = 0.0011 (6h), 0.0119 (12h), p = 0.0481 (24h), p = 1.98 x 10^-5^ (36h). **J)** Representative single-cell scatterplots of cyclin D1 and p27 protein levels colored by p-Rb status for cells treated as in (H). **K)** Schematic for bifurcation of the CDK2 activity reporter for asynchronously cycling cells computationally aligned by mitosis and analysis of CDK_high_ fraction for cells born after variable time in DMSO, LATSi, or TEADi treatment (see also Methods). **L)** Schematic representing variable time cells experienced drug relative to mitosis and 3h post-mitosis (mid-G1 indicator, left) and normalized fraction of CDK_high_ cells at 3h post-mitosis, binned by time spent in LATSi or TEADi (right). The fraction of CDK_high_ cells was normalized to the DMSO control treatment (n = 3 independent experiments, n > 5,000 cells/condition). All histogram and single-cell data (n = 8,000 cells/scatterplot) are representative of 3 independent experiments.

We confirmed that the conserved YAP target gene signature^38^ was significantly enriched after 12-36 hours of induction (Figure 3B), including the canonical YAP target genes *CYR61* and *ANKRD1* that were significantly induced within 6h (Figure 3C, S3B). We also confirmed previously-characterized induction of G1-related target genes *CDK6*^39^ and *SKP2*^40^ (Figure S3B), proliferation-supporting transcription factors (*MYBL1, ETS1, MYC,* Figure S3C)^15^, as well as TEAD and AP1 target genes^41^ (*TEAD1/2*, *FOSL1/2, JUN*, Figure S3D). Compared with the fast induction of the YAP target gene *CYR61*, we observed *CCND1* (cyclin D1) expression increasing only after 12-36 hours and only 1.4-fold. In comparison, the previously-reported suppressed gene *DDIT4*^42^ rapidly dropped whereas the expression of p27 (*CDKN1B*) and p21 (*CDKN1A*) changed only minimally over time (Figure 3D).

Since there was only a small fold change in cyclin D1 and p27 expression, we next tested whether cyclin D1 and p27 are more strongly regulated at the protein level^43,44^ following acute YAP-TEAD activation. LATS inhibitor treatment of serum-starved WT cells increased the n/c YAP ratio and we confirmed the protein-level increase of select YAP targets (CDK6 and SKP2, Figure S3E). LATS inhibition strongly increased cyclin D1 protein levels but also resulted in a small increase in p27 and transient decrease in p21 (Figure 3E, S3F). Over 3-12 hours following LATS inhibition (LATSi), the increase in the cyclin D1/p27 ratio was paralleled by increasing Rb phosphorylation (Figure 3F-G), and cells progressed to S-phase within 12-24 hours (Figure S3G). Supporting that LATS1/2 signal through YAP-TEADs, TEAD inhibition reversed the LATSi-induced changes of these protein levels and reversed the percent of proliferating cells back to control levels (Figure 3F, S3G).

We next inhibited TEAD in proliferating cells to determine how YAP-TEAD inhibition regulates cyclin D1 and p27 levels and Rb phosphorylation in G1 phase to terminate proliferation. We first confirmed that TEAD inhibition of CA-YAP cells, or of LATSi pre-treated WT cells, rapidly reduced the YAP ratio within 3 hours (Figure S3H). Markedly, however, TEAD inhibition resulted in only a delayed reduction of the nuclear level of cyclin D1 protein in G1 phase (Figure 3H). Moreover, TEAD inhibition caused a strong and slow increase of the nuclear protein levels of p27 and a relatively smaller increase in p21 levels across the population. Concordantly, the mean cyclin D1/p27 ratio and the G1 phase Rb phosphorylation both decreased but only after a delay with most cells arresting by 24-36 hours (Figure 3I, S3I). Moreover, single-cell analysis confirmed that the cyclin D1/p27 ratio still predicted whether Rb was phosphorylated in G1 as individual cells gradually reduced their cyclin D1 and increased their p27 level (Figure 3J).

To more directly measure the delay between LATS and TEAD inhibition and the decision in G1 whether to enter the next cell cycle after mitosis, we used live-cell analysis of asynchronously dividing cells expressing a reporter of CDK2 activity^45^. CDK2 activity in cycling cells bifurcates within a few hours after mitosis, distinguishing cells that enter the next cell cycle from cells that stay in G1 phase^24^. We first computationally aligned cells by the time they exited mitosis to compare subsequent cell cycle entry. We then measured the fraction of cells with high CDK2 activity at 3 hours into G1 phase and binned them by the duration of LATS or TEAD inhibition (Figure 3K).

To measure the effect of the inhibitor treatment as a function of time, we normalized the fraction of cells with high CDK2 activity in inhibitor-treated cells to the fraction in control cells to compare the relative effect of LATS or TEAD inhibition on the timing of cell cycle entry or exit. This analysis showed that LATS inhibition increased the fraction of cells activating CDK2 about 6-fold over 24 hours, while TEAD inhibition after a short delay decreased the fraction of cells about 6-fold over 24 hours (Figure 3L, S3J-K).

Thus, rather than inducing acute changes in cell cycle entry, YAP activation and inactivation regulate the cyclin D1/p27 ratio, Rb phosphorylation, and cell cycle entry gradually over a period of 24 hours. The slow kinetics of the regulation of cell cycle exit suggests that YAP is regulating the cyclin D1/p27 ratio and proliferation mostly indirectly. This raises the question of what mechanism mediates this slow acting component of YAP-regulated cell cycle control.

### YAP activation induces and prolongs proliferation by increasing EGFR signaling

Our RNAseq time course analysis supported earlier studies that YAP activation increases multiple factors related to receptor signaling^16,46–48^ (Figure 4A, S4A). These factors included receptors and secreted ligands known to regulate cell proliferation including *ERBB3* and *AXL* receptors^3^, along with ligands like *FGF1/2* and the EGFR ligand precursor *HBEGF* (paralog of *AREG*)^35,49^, providing a potential cell autonomous activation of EGFR and other receptors. We were intrigued by this possibility since the slow kinetics of the expression changes matched the delay in the changes of the cyclin D1/p27 ratio in the G1 phase after TEAD inhibition.

**Figure 4.**
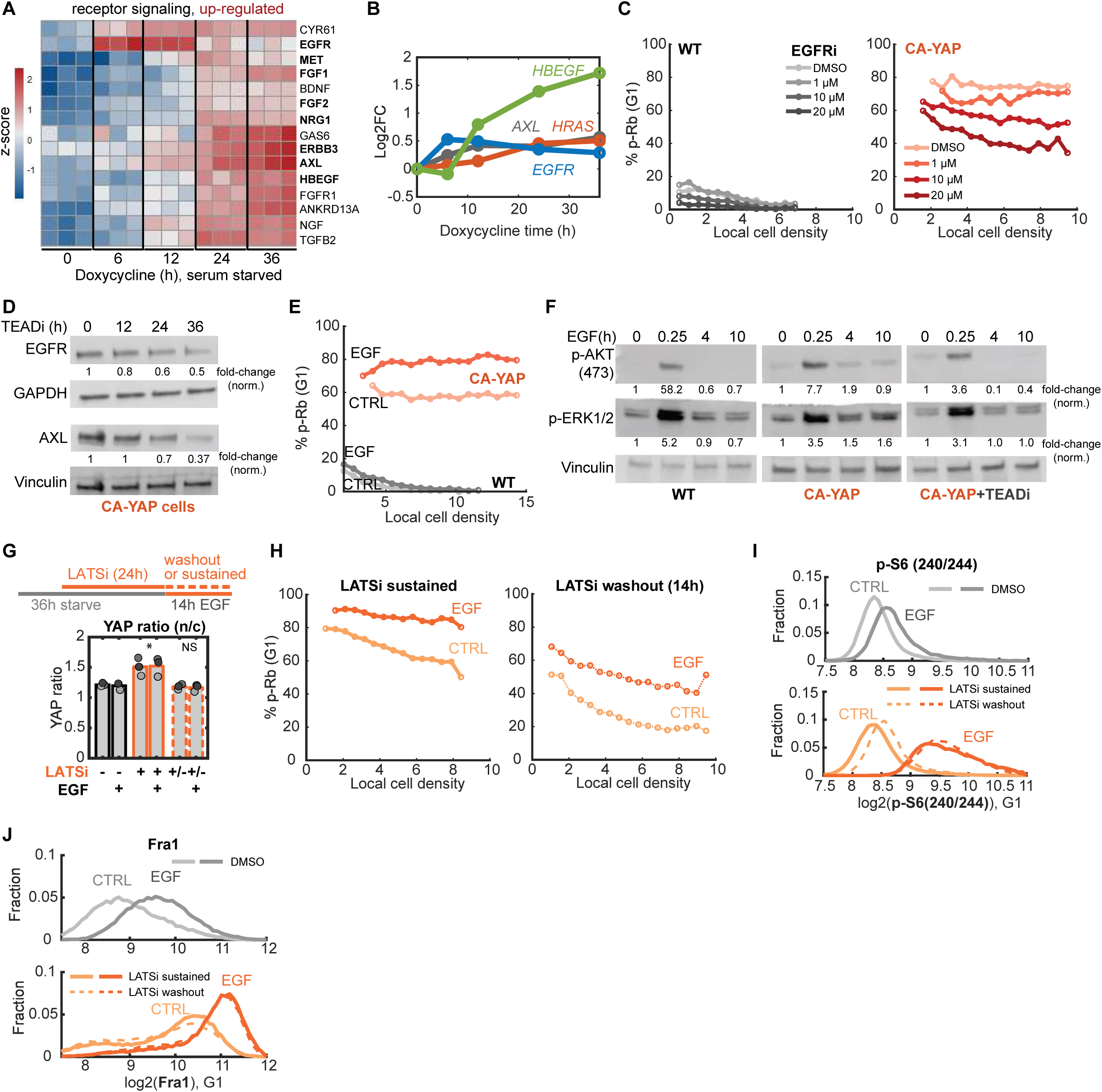
YAP activation induces and prolongs proliferation by increasing EGFR signaling. **A)** Heatmap representation of differential expression (row z-score) of select up-regulated ligand and receptor genes, grouped by 3 independent replicates and time points. **B)** Log2 fold-change (Log2FC) of select receptor signaling-related genes, *AXL* (direct YAP target, grey), *EGFR* (blue), *HBEGF* (green), and *HRAS* (red). Adjusted p-values: *AXL,* 6.20 x10^-4^ (6h), 9.11×10^-12^ (12h), 2.33×10^-11^(24h), 2.19×10^-21^ (36h); *EGFR,* 4.97×10^-16^ (6h), 1.07×10^-14^ (12h), 1.07×10^-8^ (24h), 1.23×10^-5^ (36h); *HBEGF*, 3.90×10^-4^ (12h), 8.57×10^-12^ (24h), 5.24×10^-18^ (36h); *HRAS*, 4.10×10^-4^ (24h), 6.05×10^-5^ (36h). **C)** Percent p-Rb binned by local cell density for WT (left) and CA-YAP (right) cells serum-starved and treated with gefitinib (EGFRi, 0-20 μM) for 24h. See also **Figure S4.** **D)** Western blot analysis of EGFR and AXL from whole-cell lysates of serum-starved CA-YAP cells treated with TEADi for 0-36h, with GAPDH and Vinculin loading controls. Values indicate fold-change difference relative to 0h control (after normalization to loading control). Data are representative of 2 independent experiments. **E)** Percent p-Rb binned by local density for serum-starved WT and CA-YAP cells after 14h EGF (25 ng/mL). Data are representative of 2 independent experiments, n > 25,000 cells/condition. **F)** Western blot analysis of AKT and ERK phosphorylation levels in whole-cell lysates of serum-starved WT (left), CA-YAP (middle), and TEADi pre-treated CA-YAP cells (right) after 0-10h EGF treatment (25 ng/mL). Values indicate fold-change difference relative to 0h control (after normalization to loading control). **G)** <u>Top</u>: experimental conditions for serum-starvation of WT cells, pre-treatment with LATSi, and sustained or washout of LATSi during 14h EGF treatment (25 ng/mL). <u>Bottom</u>: Mean YAP ratio (n/c) for treatment conditions. N = 3 independent experiments, matched colors indicate conditions from the same experimental replicate. Student’s t-test to control, YAP ratio: LATSi sustained, p= 0.0278 (CTRL), 0.027 (EGF); LATSi washout, not significant (NS). **H)** Percent p-Rb binned by local cell density for cells with LATSi sustained (left) or washout (right) with or without EGF treatment, n > 20,000 cells/condition. **I-J)** Histogram distribution of phosphorylated-S6 (p-S6 240/244, I) and Fra1 protein levels for DMSO-treated (<u>top</u>, CTRL) or LATSi-pretreated cells (<u>bottom</u>, LATSi sustained and washout). Data are representative of 3 independent experiments. All single-cell data represent values from G1-gated cells.

Since expression levels of *AXL* (a direct YAP target gene)^15,50,51^, *EGFR,* and *HRAS* were all gradually increased by YAP activation, along with the EGFR-activating ligand *HBEGF* (Figure 4B), we first considered whether the residual proliferation observed in serum starved cells at low local density might be induced by YAP through activation of EGFR signaling. We observed that the small portion of WT cells proliferating in the absence of mitogens could be inhibited by titration of the EGFR inhibitor gefitinib, even in the absence of exogenous EGF (Figure 4C). This EGFR inhibition mirrored the TEAD inhibition (Figure S1F), supporting that YAP-TEAD activity may signal through EGFR at low stimulation conditions to promote proliferation. Nevertheless, EGFR inhibition in CA-YAP cells could only partially suppress Rb phosphorylation (Figure 4E, Figure S4B), suggesting that additional receptor signaling pathways are contributing to the YAP-induced proliferation.

We next determined how rapidly TEAD inhibition changes the protein level of EGFR relative to its established target AXL. Western blot analysis revealed that their total levels gradually decreased after TEAD inhibition in CA-YAP cells over a similar time scale as the loss of proliferation (Figure 4D). Since EGFR protein levels were YAP-dependent, we further tested whether YAP activation increases EGF sensitivity. Indeed, while EGF stimulation caused only a small increase in Rb phosphorylation in WT cells, Rb phosphorylation was much more strongly increased in CA-YAP cells (Figure 4E). Furthermore, while both WT and CA-YAP cells had similarly low basal AKT and ERK phosphorylation, the CA-YAP cells kept their phosphorylated AKT and ERK levels higher for at least 10 hours after EGF stimulation compared to WT cells, again consistent with sensitized EGF signaling (Figure 4F). As a control, we also confirmed that TEAD inhibition of CA-YAP cells prevented this prolonged AKT and ERK signaling.

Finally, we tested whether there is an extended period of enhanced EGF signaling also after transient LATS inactivation upstream of YAP. We used WT cells pre-treated with LATSi and then either maintained (sustained) or washed out the LATSi prior to a 14-hour addition of EGF (Figure 4G). While LATSi washout reduced the n/c YAP ratio and nuclear YAP levels back to control conditions (Figure 4G, S4C), cells retained sensitivity to EGF for at least 14 hours, as evidenced by the increased p-Rb (Figure 4H, Figure S4D), S6 phosphorylation (p-S6, a readout of PI3K/mTOR activity, Figure 4I, S4E), and nuclear Fra1 levels (a YAP target but also a readout of integrated MEK-ERK activity^52^, Figure 4J). In support of a general prolongation of proliferation signaling, we also found that transient LATSi treatment sustained the fraction of G1 cells with phosphorylated Rb after HGF or lysophosphatidic acid (LPA) stimulation (Figure S4F).

We conclude that YAP activity increases the activity of EGFR in the absence of added EGF in an autocrine manner^16,49^, while also increasing the sensitivity of cells to exogenous EGF stimulation. This sensitization extends to other receptors since gefitinib could only suppress a fraction of the YAP-induced proliferation levels and LATS-inhibited cells were more proliferative following treatment with other mitogens. Notably, we showed that increased EGFR signaling persisted for over 14 hours after YAP inactivation, suggesting that YAP is inducing a prolonged proliferation response since its inactivation only terminates proliferation and signaling after a delay.

### YAP activation signals primarily through MEK-ERK to increase the cyclin D1/p27 ratio

We next examined which signaling pathways downstream of the YAP-induced receptor activation control the cyclin D1/p27 ratio. We focused on PI3K-AKT-mTOR and MEK-ERK signaling^46,53^, since various functions of YAP activation are mediated through mTOR activation,^54,55^ and both mTOR and MEK-ERK can regulate cyclin D1 and p27 levels^18,33,56^ (Figure 5A). We measured phosphorylation of AKT and S6 protein (p-S6) to monitor PI3K pathway activity and protein levels of the transcription factor Fra1 to monitor the ERK pathway. Serum-starved CA-YAP cells displayed both elevated p-AKT(S473) and p-S6(240/244) as well as Fra1 levels (Figure 5B-C), which similarly depended on YAP-TEAD activity, as TEAD inhibition reduced their phosphorylation and protein levels in CA-YAP cells (Figure 5D).

**Figure 5.**
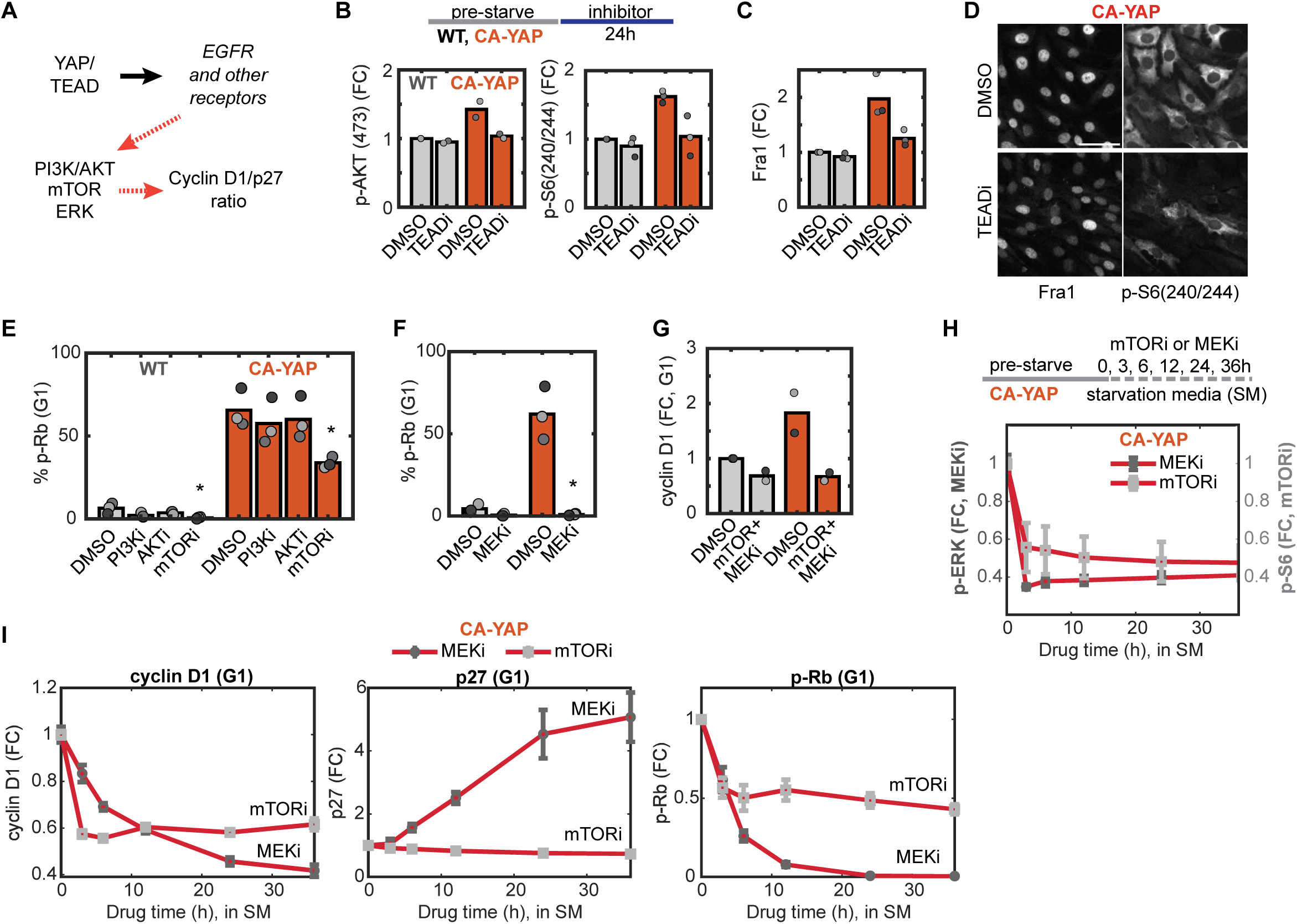
YAP activation signals through ERK and mTOR to increase the cyclin D1/p27 ratio. **A)** Schematic of proposed indirect regulation of cyclin D1/p27 ratio by YAP/TEAD via EGFR-activated signaling pathways. **B-C)** <u>Top</u>: experimental conditions for serum-starvation of WT and CA-YAP cells followed by 24h treatment with TEAD inhibitor for 24h in starvation media. <u>Bottom</u>: Mean fold-change (FC) in protein levels of p-AKT(473, left) and p-S6(240/244, right) (B) and Fra1 (C) for serum-starved WT and CA-YAP cells treated with TEADi for 24h (normalized to WT/DMSO condition). N = 2 (p-AKT) or n = 3 (p-S6, Fra1) independent experiments. **D)** Representative multiplexed images for Fra1 and p-S6(240/244) staining in CA-YAP cells treated with TEADi as in (B). Scale bar = 50 μm. **E-F)** Mean percentage p-Rb for WT and CA-YAP cells treated with PI3K (LY294002, 10 μM), AKT (MK2206, 200 nM), or mTORC1/2 (Torin2, 100 nM) inhibitors (E) or MEK (PD0325901, 100 nM) inhibitor (F) as in (B). N = 3 independent experiments. Student’s t-test: WT/mTORi, p = 0.0372; CA-YAP/mTORi, p = 0.0103 (E). CA-YAP/MEKi, p = 0.0028 (F). See also **Figure S5**. **G)** Mean FC in cyclin D1 protein levels for WT and CA-YAP cells treated with combined mTORi and MEKi relative to WT/DMSO condition as in (B). N = 2 independent experiments. **H)** <u>Top</u>: experiment conditions for mTORi or MEKi time course treatment of serum-starved CA-YAP cells. <u>Bottom</u>: fold-change in p-ERK (MEKi-treated, dark grey) and p-S6(240/244, mTORi-treated, light grey) following drug treatment. Mean ± SEM (n = 2 independent experiments, 2-3 replicate wells/experiment). **I)** Fold-change in protein levels of cyclin D1, p27, and p-Rb for MEKi and mTORi for CA-YAP cells treated as in (H). Mean ± SEM.

To assess the functional importance of PI3K-AKT-mTOR signaling, we inhibited PI3K (LY-294002) or AKT (MK2206), which had only small inhibitory effects on Rb phosphorylation in CA-YAP cells (Figure 5E), despite reducing the p-AKT(S473) and p-S6(240/244 and 235/236) levels (Figure S5A-B). Nevertheless, inhibiting mTORC1/2 (mTORi, Torin2) caused an approximately 50% reduction in proliferation (Figure 5E). In contrast, inhibition of the obligate upstream regulator of ERK, MEK (MEKi, PD0325901), completely suppressed Rb phosphorylation and readouts of ERK activity (Figure 5F, S5C-D).

In support of YAP regulating cyclin D1 protein levels in G1 phase through mTOR and MEK/ERK, rather than doing so directly, MEK inhibitor alone (Figure S5D) or the combined inhibition of MEK and mTOR fully reduced the cyclin D1 level in CA-YAP cells to the level of WT cells (Figure 5G, S5E). A time course analysis showed that the drop in cyclin D1 levels is rapid after mTOR inhibition but slower and stronger after MEK inhibition even though both inhibitors rapidly suppressed p-S6 and p-ERK, respectively (Figure 5H-I). As mTOR can rapidly control translation of cyclin D1 mRNA^57^, this fast regulation was anticipated. In contrast, mTOR inhibition did not alter p27 levels but MEK inhibition not only reduced cyclin D1 levels but also slowly and strongly increased p27 levels, explaining why MEK inhibition suppresses YAP-dependent Rb phosphorylation more completely than the regulation of mTOR. Thus, transient YAP-mediated proliferation responses result from an increase and delayed decrease of the activity of EGFR and other receptors, which in turn transiently increases the cyclin D1/p27 ratio through reversibly regulating MEK-ERK and, to a lesser degree, mTOR signaling pathways.

### Partial activation of YAP from ablating Merlin can be reversed by increased contact inhibition

Previous studies suggested that EGFR signaling and contact inhibition of proliferation are also regulated independently of YAP through regulation of an upstream Hippo pathway activator, Merlin/NF2^58–60^. We were particularly intrigued to study Merlin since its genetic loss increases tissue size but with conflicting reports on the relevance of YAP signaling versus mitogen responses. We generated clonal Merlin knockout RPE1 cells (Merlin KO) using CRISPR/Cas9 and confirmed that these cells had increased nuclear localization of YAP that could be suppressed by TEAD inhibition (Figure S6A-B).

Strikingly, unlike CA-YAP, Merlin KO increased proliferation without abolishing contact inhibition of proliferation. Instead, a higher cell density was required to inhibit proliferation compared to WT, both at low and high serum levels (Figure 6A, S6C). Moreover, inhibiting TEAD in Merlin KO cell suppressed Rb phosphorylation in both cases to levels similar as TEAD-inhibited WT cells (Figure S1F), supporting a model whereby Merlin regulates proliferation primarily through YAP-TEAD activity^9^. Crucially, YAP activity in Merlin KO cells could be as effectively inhibited by contact inhibition as in WT cells but required a higher local cell density to reach the same YAP inhibition (both at low and high serum levels, Figure 6B).

**Figure 6.**
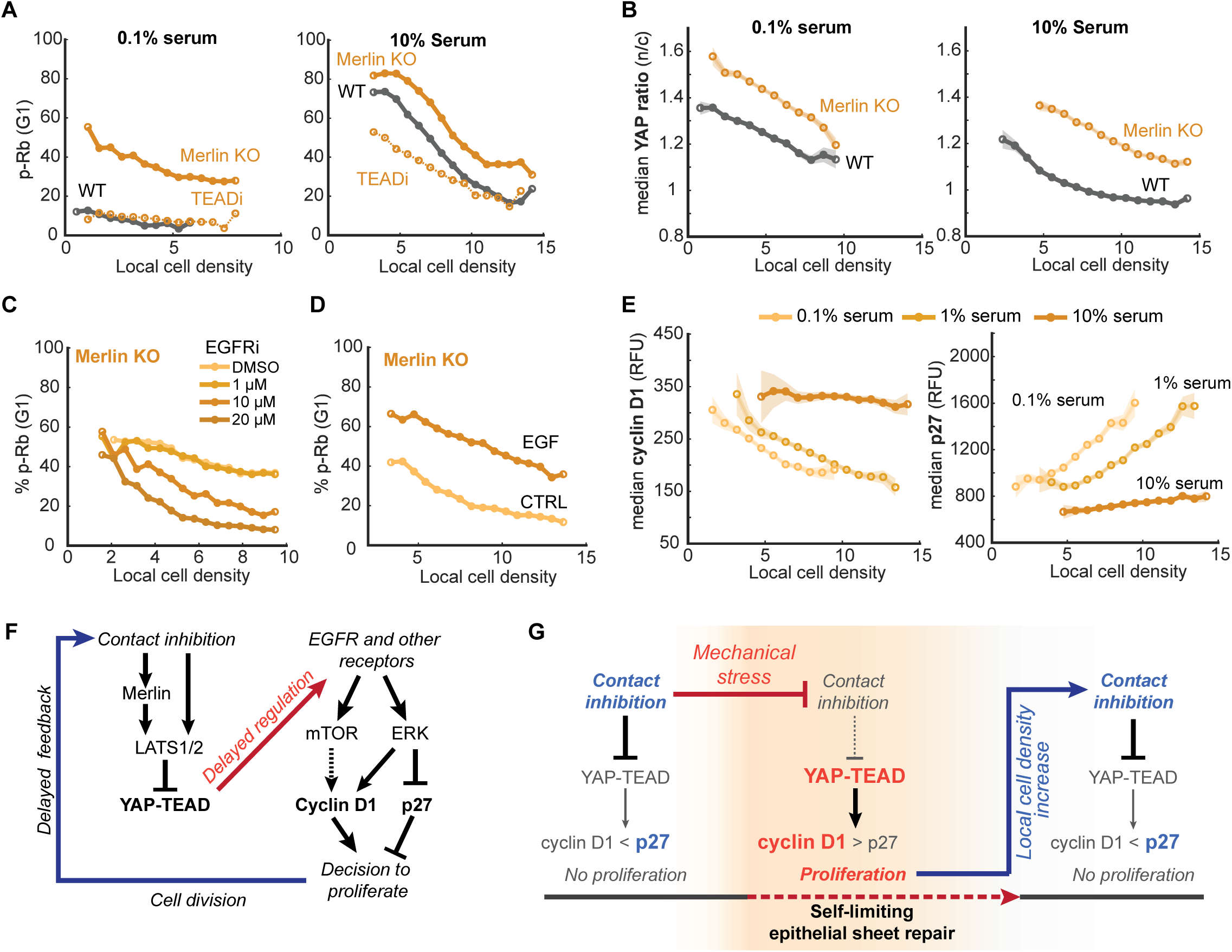
Partial activation of YAP by Merlin knockout can be reversed by increased contact inhibition. **A)** TEAD-dependent hyperproliferation of Merlin KO cells. Percent p-Rb binned by local density for Merlin knockout cells (Merlin KO, yellow) in 0.1% (left) or 10% serum (right) treated with DMSO or TEADi (0.5 μM) compared with WT cells (grey). **B)** Median YAP ratio (n/c) binned by local cell density for Merlin KO cells in 0.1% (left) or 10% serum (right), compared with WT cells (same experiment as in Figure 1E). Data are representative of 3 independent experiments. Shaded error are 95% confidence intervals. **C)** Percent p-Rb binned by local density for serum-starved Merlin KO cells treated with gefitinib (EGFRi). Data are representative of 2 independent experiments. **D)** Percent p-Rb binned by local cell density for serum-starved Merlin KO cells with control (CTRL) or EGF (25 ng/mL) treatment for 12h. Data are representative of 2 independent experiments. **E)** Median cyclin D1 (left) or p27 (right) protein levels for Merlin KO cells in titrated serum. Shaded error are 95% confidence intervals. Data are representative of 2 independent experiments. **F)** Model for Hippo-pathway inhibition of YAP-TEAD activity, which regulates signaling through and downstream of EGFR to control the cyclin D1/p27 ratio. **G)** Model proposing transitions between low and high YAP-TEAD activity as contact inhibition is temporarily reduced then recovered after a delay during epithelial sheet regeneration. For all representative binned plots, data are pooled from at least 2 wells, n > 100 cells/bin, n > 10,000 cells/plot. All quantifications are for G1-gated cells. See also **Figure S6**.

We next examined the role of EGFR in the Merlin KO cells, as a possible mechanism mediating serum-independent proliferation downstream of the higher YAP signaling. Like CA-YAP cells, Merlin KO cells displayed activated EGFR signaling in the absence of added EGF, since the EGFR inhibitor gefitinib partially reduced the percentage of proliferating cells (Figure 6C, S6D). Like the transiently LATS-inhibited cells, the Merlin KO cells had increased sensitivity to EGF while still retaining a strong local cell density dependence (Figure 6D). In addition, we found that Merlin KO cells increased proliferation in response to other receptor tyrosine kinase ligands (HGF and FGF, Figure S6E), again consistent with YAP inducing a broad upregulation of receptor sensitivity that is not limited to EGFR.

Finally, we found that Merlin KO cells still suppressed cyclin D1 and increased p27 levels at higher local cell density (Figure 6E, S6F), demonstrating that these cells suppress proliferation through the same mechanism as WT cells: inhibiting YAP activity and downregulating mitogen signaling and the cyclin D1/p27 ratio. However, Merlin KO cells employ alternative mechanisms from inactivating Merlin^61,62^ to contact inhibit YAP.

We conclude that even though constitutive YAP activation abolished contact inhibition of proliferation, cells can robustly contact inhibit YAP-TEAD even if Hippo signaling is reduced and YAP activation increased. These studies also support that Merlin signals through YAP to control EGFR and other receptor signals that together control the cyclin D1/p27 ratio. However, cells that have such an induced YAP activity now require higher local cell density to suppress YAP activity and proliferation.

## DISCUSSION

Our study shows that contact inhibition of YAP and mitogen signaling control the cyclin D1/p27 ratio and cell cycle entry in G1 phase through a balance mechanism that shapes an intrinsically transient YAP-induced proliferation response (Figure 6F). Our study supports a model where YAP activation controls proliferation by gradually increasing the expression of HB-EGF and other RTK ligands, but also various growth factor receptors and known regulators of mitogen signaling^3,54,63^ that mediate signal sensitivity and duration. This regulation explains how YAP activation is increasing EGF and other mitogen signals in a cell autonomous manner (Figure 4C), while simultaneously sensitizing cells to exogenous stimulation by EGF and other mitogens (Figure S4F, S6E), which is likely relevant for the role of YAP activation in the inflammatory injury response^64,65^. We further show that YAP activation increases downstream signaling through MEK-ERK signaling and, to an unexpectedly lesser degree, mTOR signaling, to control the nuclear protein levels of cyclin D1 and p27 (Figure 5I). Most importantly, our study reveals a predominant role of both cyclin D1 and p27 in YAP-induced proliferation. We showed that YAP-stimulated cells decide in the G1 phase whether to enter the cell cycle by increasing the nuclear cyclin D1/p27 ratio above a critical activity threshold, or to remain quiescent or exit the cell cycle by staying below this threshold (Figures 2D-E, 3G,J).

Moreover, as the YAP-induced cell division increases the local cell density, we show that the ensuing contact inhibition of YAP-TEAD causes a delayed inhibition of mitogen signaling, which lowers the cyclin D1/p27 ratio to terminate the proliferation response. Thus, there are two delay mechanisms ensuring that the proliferation response is prolonged: cells wait to inhibit YAP until cell division has increased the local cell density, then the resulting contact inhibition of YAP lowers the cyclin D1/p27 ratio and arrests proliferation after a delay (Figure 6G). YAP signals often fluctuate^66–68^, and the slow integration of YAP signals further ensures that the proliferation response is robustly started and ended. Slow integration provides a filter mechanism which prevents short-lived YAP activity changes from triggering unwanted cell cycle entry or premature cell cycle exit.

We also evaluated the robustness of YAP inactivation by using ablation of Merlin/NF2, a suppressor of Hippo signaling to YAP^69–71^. While Merlin has been shown to have additional signaling functions at the plasma membrane, our study shows that the increased EGFR signaling and proliferation in Merlin KO cells can be reversed by TEAD inhibition (Figure 6A), suggesting that Merlin primarily signals through YAP-TEAD to control EGFR signaling, the cyclin D1/p27 ratio, and proliferation. In support that the termination of YAP signaling and proliferation is robust, our study shows that contact inhibition is merely shifted but not abolished when YAP activity is increased by ablation of Merlin, and cells can still effectively terminate proliferation but at higher local cell density. Moreover, since the loss of Merlin can boost proliferation without a loss of contact inhibition, this result further suggests that Merlin has a unique advantage as a drug target, since a potential Merlin inhibitor would trigger a self-limiting proliferation response. This could offer a safer alternative strategy to LATS inhibition, which was recently shown to cause a widespread and uncontrolled proliferation response and de-differentiation in mice^6^.

We also found that inhibiting MEK-ERK, rather than mTOR, in CA-YAP cells greatly reduces the cyclin D/p27 ratio and stops proliferation. This crucial role of MEK-ERK in YAP-induced proliferation was surprising, since oncogenic YAP mutants can drive resistance to KRAS inhibition and KRAS primarily regulates proliferation through MEK-ERK and mTOR activation^28,46,72^. The resistance to KRAS inhibitor may therefore by mediated by YAP-mediated direct regulation of cell cycle entry. We confirmed that YAP activation can also directly induce cyclin D1 as well as SKP2, which can degrade p27^73,74^. Indeed, we found that this upregulation may be particularly important at high levels of YAP activation^75,76^, since DOX induction of CA-YAP in high serum conditions can still increase cyclin D1 and proliferation when MEK is inhibited (Figure S6G-H).

### Concluding remarks

YAP activation must be transient to drive the epithelial regenerative response without excessive hyperplasia. Our study identifies a delayed, self-limiting control mechanism that allows YAP to trigger such a commensurate proliferation response. We show that transient proliferation responses are initiated by YAP activation increasing EGFR and mitogen signaling to increase the cyclin D1/p27 ratio above a critical threshold. The delayed contact inhibition-mediated inactivation of YAP then terminates proliferation by gradually lowering mitogen signaling and the cyclin D1/p27 ratio below this critical threshold (Figure 6G). In contrast, we demonstrate that persistent inactivation of LATS1/2 or activation of YAP activity abolishes contact inhibition and promotes persistent mitogen signaling that keeps the cyclin D1/p27 ratio high, explaining how oncogenic mutations that constitutively inactivate LATS1/2 or activate YAP elicit unrestricted proliferation.

## Acknowledgments

We thank Nathaniel Kastan and James Hudspeth for early access to a previous iteration of the LATS1/2 inhibitor (TRULI); Alexander Pfannenstein, Marielle Koeberlin, and Sanjeev Sharma for technical support; the Stanford Shared FACS Facility and Weill Cornell FACS Facility for cell sorting; the Weill Cornell Genomics Core for library prep and RNA-sequencing; Marielle Köberlin for critical reading of the manuscript; and James Ferrell, Alexander Dunn, Daniel Jarosz, Ovijit Chaudhuri, Elizabeth Kiddie, members of the Meyer and Teruel laboratories for technical feedback and helpful discussions. This work was funded by an NIH T32 training grant (5 T32 GM 113854-3), NIH grants T.M. (R35 GM12702601) and M.N.T. (R01 DK131432); K.R.F. was supported by an NSF Graduate Research Fellowship (DGE-1656518).

## Author contributions

Conceptualization, K.R.F., Y.F., and T.M.; Methodology, K.R.F., N.R., Y.F., and T.M.; Experiments, visualization, and analysis, K.R.F.; Manuscript preparation, K.R.F., M.N.T., and T.M.; Funding acquisition, T.M., and M.N.T.

## Resource availability

Immunofluorescence protocols, and cell lines are available upon request to the lead contact. Plasmids generated in this study will be deposited to Addgene.

## Data and code availability

MATLAB image-processing scripts and iterative imaging analysis pipeline are previously published^22,77^ and publicly available on Github (https://github.com/MeyerLab/image-analysis-ratnayeke-2022).

Additional information is available upon request from the lead contact.

## METHODS

### Cell culture

All experiments were performed with human retinal pigmented epithelial cells (RPE1-hTERT immortalized, ATCC Cat# CRL-4000) unless specified otherwise. RPE1 were cultured in DMEM/F12 growth media with HEPES (Thermo Scientific Cat#11039047), supplemented with 10% fetal bovine serum (Millipore Sigma Cat#F4135-500ML) and passaged using trypsin-EDTA (0.05%, Gibco Cat# 25300054). Human mammary epithelial cells (MCF10A, ATCC Cat# CRL-10317, RRID:CVCL_0598) were cultured in DMEM/F12 growth media with HEPES supplemented with 5% horse serum (Gibco Cat# 16050122), 20 ng/mL EGF (PeproTech Cat# C8052), 0.5 μg/mL hydrocortisone (Sigma Cat#H0888), 100 ng/mL cholera toxin (Sigma Cat# C8052), and 10 μg/mL insulin (Sigma Cat#I1882). For serum starvation experiments, cells were first washed with serum-free DMEM/F12 media, then maintained in serum-free (0%) or 0.1% serum-containing DMEM/F12 (serum-starvation conditions). 0.1% serum-containing media had similar proliferation as 0% serum-containing media, but cells were slightly more adherent to glass-bottom plates for subsequent immunofluorescence protocols and therefore used for a subset of experiments. For mitogen-release experiments, mitogens and growth factors (EGF, HGF, and FGF at 25 ng/mL and LPA at 1 μM) were diluted in starvation media. HEK-293T cells used for lentivirus production were cultured in DMEM growth media (Gibco Cat# 11995065) supplemented with 10% fetal bovine serum. Cells were cultured at 37 °C and 5% CO2.

### Plasmid generation

pSpCas9(BB)-2A-Puro (PX459) V2.0 was a gift from Feng Zhang (Addgene # 62988, RRID:Addgene_62988). Phosphorylated and annealed sgRNA oligos against *NF2* (see Cell line generation below) were cloned into restriction enzyme-digested PX459^78^.

pQCXIH-Myc-YAP-5SA was a gift from Kun-Liang Guan (Addgene #33093, RRID: Addgene_33093). Doxycycline-inducible, myc-epitope-tagged CA-YAP (YAP5SA) was generated using Gibson assembly of PCR-amplified myc-YAP5SA with restriction enzyme-digested plasmid backbone derived from a bicistronic vector with a TetOn promoter driving target gene expression and PGK-driven rtTA expression, pCW-Cas9 (Addgene #50661, RRID: Addgene_50661, a gift from Eric Lander and David Sabatini).

### Cell line generation

Merlin knockout cells were generated by transient transfection of parental RPE1 cells with pX459 (2 ng/uL, containing gRNA scaffolds targeting Merlin/Nf2) and Lipofectamine 2000 (1:500, Thermo Cat# 11668019) in Opti-MEM (Fisher Scientific Cat#31-985-070) and growth media overnight, then exchanged for growth media with puromycin selection (10 μg/mL, Invivogen Cat#ant-pr-1) for 72h. After a recovery period in selection-free media, cells were diluted to 0.5 cells/100 μL and seeded to 96-well plates for single-cell clonal expansion. 24 single-cell clones were further expanded for preliminary knockout assessment by western blot for loss of Merlin protein expression. 2 clones (Merlin KO-clone 1 forward: CACCGCACAGTGGCCTGGCTCAAAA, reverse: AAACTTTTGAGCCAGGCCACTGTGC; Merlin KO-clone 2 forward: CACCGTGATTTGGTGTGCCGGACTC, reverse: AAACGAGTCCGGCACACCAAATCAC) were selected for validation by genomic DNA purification (Qiagen CAT#69504), PCR of the Merlin locus and sequencing. After initial characterizations confirming Merlin KO-clone 1 and Merlin KO-clone 2 were similarly proliferative (data not shown), we proceeded to use KO-clone 1 for all subsequent experiments in this study.

Lenti-EF1a-dCas9-VPR-Puro (Addgene # 99373, RRID: Addgene_99373) was a gift from Kristen Brennard and stably expressed in RPE1 cells using puromycin selection (10 ug/mL, Invivogen Cat#ant-pr-1). CA-YAP and dCas9-expressing cells were generated using third-generation lentiviral transduction. Lentivirus was generated in HEK-293T cells by transfection with packaging and envelope plasmids pMDLg/pRRE (Addgene # 12251, RRID: Addgene_12251), pRSV-rev (Addgene #1225, RRID: Addgene_1225), and pCMV-VSV-G (Addgene #8454, RRID: Addgene_8454) in Lipofectamine 2000 and Opti-MEM. After 72h transfection, supernatant was passed through a 0.22 uM filter (Millipore Cat# SCGP00525) and concentrated using 100 kDa centrifugal filters (Millipore Cat# UFC910024). Lentivirus was then frozen or directly transduced into cells. 72h after lentiviral transduction, cells were split into antibiotic-selection media or, if expressing a fluorescent construct, expanded prior to sorting by a BD Influx cell sorter (Stanford Shared FACS Facility, Weill Cornell FACS Facility). RPE1 reporter cell lines were generated by transduction with pLV-H2B-miRFP670 (nuclear marker), CSII-pEF1a-hDHB(994-1087)-mTurquoise (cyclin E/A-CDK reporter), and the FUCCI cell-cycle reporters CSII-pEF1a-mVenus-hGeminin(1-110) (APC/C^Cdh1^ reporter) and pLV-mCherry-hCDT1(1-100)ΔCy (S-phase start^79^), as described previously^77^.

### siRNA, sgRNA, and plasmid transfection

All oligo sequences are listed in Table S1. Pooled siRNA (ON-TARGETplus, Horizon Discovery Biosci.) were used for siCtrl (Non-targeting Control, Horizon Cat#D-001810-10-05), siCycD1 (siCCND1, Horizon Cat# L-003210-00-0005), and sip27 (siCDKN1B, Horizon Cat# L-040178-00-0005). For CRISPRa-mediated overexpression of genes through targeting of endogenous promoters, pooled sgRNA (Horizon) were used for sgCtrl (non-targeting control, Horizon Cat# U-009500-10-05), sgCycD1 (sgCCND1, Horizon Cat# P-003210-01-0005), and sgp27 (sgCDKN1B, Horizon Cat# P-003472-01-0005) with tracRNA (Edit-R tracrRNA, Cat# U-002005-05). RPE1 cells were transfected with siRNA and sgRNA using DharmaFECT 1 (1:500 in Opti-MEM, Dharmacon Cat# T-2001-03) according to manufacturer protocol using 20 nM siRNA, or sgRNA and tracrRNA (in 1:1 ratio, according to manufacturer protocol). Cells were incubated in transfection mixture for 6-12h in growth media, then replaced with media appropriate for the experiment conditions and assessed 24-48h after transfection.

### Drugs and mitogens

Human growth factors and mitogens were dissolved in sterile, ultrapure water (Fisher Scientific Cat#10977023) and used at the following concentrations unless otherwise stated: 25 ng/mL recombinant EGF (PeproTech cat#AF-100-15), 25 ng/mL recombinant HGF (Life Technologies Cat#PHG0254), 25 ng/mL recombinant FGF basic (R&D Systems Cat#233-FB-025), 1 uM LPA (oleoyl-L-alpha-lysophosphatidic acid, Sigma-Aldrich Cat#L7260-1MG).

Drugs were dissolved in DMSO (Sigma-Aldrich Cat# D2650) and used at the following concentrations unless otherwise stated: 0.5 μM TDI-011536 (Selleck Chemicals Cat#E1314, CAS#2687970-96-1, LATSi), 5 μM TRULI (a gift from James Hudspeth, CAS#1424635-83-5), 1 μM GNE-7887 (ChemieTek Cat#CT-GNE7883, CAS# 2648450-42-2,TEADi), 10 μM gefitinib (Selleck Chemicals Cat#S1025, CAS#184475-35-2, EGFRi), 10 μM LY294002 (Cayman Chemical Cat#70920, CAS#154447-36-6, PI3Ki), 200 nM MK2206 (Cayman Chemical Cat#11593, CAS#1032350-13-2, AKTi), 100 nM Torin2 (Cayman Chemical Cat#14185, CAS#1223001-51-1, mTORi), 100 nM PD0325901 (Selleck Chemicals Cat#S1036, CAS#391210-10-9, MEKi), 1 μM palbociclib (PD-0332991, Selleck Chemicals cat#S1116, CAS#827022-32-2, CDK4/6i), 0.5 ug/mL doxycycline hyclate (Sigma-Aldrich cat#D9891, CAS#24390-14-5, DOX), 10 μM 5-ethynyl-2’-deoxyuridine (EdU, Cayman Chemical Cat#20518, CAS# 61135-33-9). For drug treatments and time courses, control conditions were treated with DMSO, except mitogen release controls (mock), which were dissolved in water. For all EdU-incorporation, cells received an EdU spike-in (10 µM, in serum-free media) for 12-15 min prior to fixation.

### Western blotting

Cells grown in 6-well plates were washed once in cold, sterile PBS and lysed in chilled, 1x RIPA buffer (Boston BioProducts Cat#BP-115-5X) or non-denaturing lysis buffer (20 mM HEPES, 150 mM NaCl, 2 mM EDTA, 0.1% IGEPAL-CA630) containing HALT protease and phosphatase inhibitor cocktail (1:100, Thermo Fisher Cat#78439). Lysates were harvested on ice using cell scrapers, sheared by syringe (25G) and centrifuged at 15,000 RCF for 15 min at 4C. Supernatant was stored at −20C. Protein concentration was calculated by Pierce BCA Protein Assay (Thermo Fisher Cat#23225) according to manufacturer protocol and a BSA standard curve.

10-30 ug protein was loaded onto 7.5%, 12%, or 4-20% Mini-PROTEAN TGX gels for resolution by SDS-PAGE in 1x Laemmli sample buffer (Fisher Scientific, Cat#AAJ60660AC, or Bio-Rad Cat#1610737) alongside Page RulerPlus (Thermo Scientific Cat#26620) molecular ladders in Tris/Glycine/SDS running buffer (Bio-Rad Cat#161-0772). Gels were run at 150V for 1h then transferred to 0.2 um PVDF membranes by semi-dry transfer (Bio-Rad Trans-Blot SD, Cat# 1703940) in Tris/Glycine buffer (Bio-Rad Cat# 1610734) with 10% methanol, or semi-dry transfer (Bio-Rad Trans-Blot Turbo Cat# 1704150) with Trans-Blot Turbo Transfer Packs (Bio-Rad Cat# 1704156). Membranes were washed in TBST (20 mM Tris, pH 7.5, 150 mM NaCl, 0.1% Tween-20), blocked for 30 min in 5% milk/TBST (VWR Cat#10128-602), and incubated at RT for 3h or overnight at 4C in primary antibody dilutions in 5% BSA/TBST + 0.01% NaN3. Primary antibodies used were rabbit anti-AXL (1:1000, Cell Signaling Technology Cat# 8661, RRID:AB_11217435), rabbit anti-EGFR (1:1000, Cell Signaling Technology Cat# 4267, RRID:AB_2246311), rabbit anti-GAPDH (1:1000, Cell Signaling Technology Cat# 5174, RRID:AB_10622025), mouse anti-Vinculin (1:1000, Thermo Fisher Scientific Cat# 14-9777-80, RRID:AB_2573027), rabbit anti-Merlin (1:1000, Cell Signaling Technology Cat# 12888, RRID:AB_2650551) (see also Table S2).

After 2x washes in TBST, membranes were incubated with anti-rabbit or anti-mouse HRP-conjugated secondary antibodies (1:5000, CST#7074, RRID: AB_2099233 or CST#7076, RRID: AB_330924) or fluorophore-conjugated secondary antibodies (1:10,000, LI-COR Biosciences, IRDye 680 LT CAT#925-68020, or IRDye 800CW, CAT#925-32210) in 5% BSA/TBST for 1h at RT. Membranes treated with enhanced chemiluminescence substrate (Thermo Fisher Cat#PI34080) were detected by film (Thomas Sci. Cat#EK-5130) or Licor Odyssey Fc. Membranes treated with fluorophore-conjugated secondary antibodies were imaged on a Licor Odyssey Fc. Normalized protein level changes were calculated using FIJI ^80^ by measuring the integrated intensity of a set box size encompassing the protein band. The background signal was measured by applying the same box to an adjacent region of the blot and subtracted from the median protein intensity. An equivalent box size was used for reference/loading control protein bands as the protein of interest.

### Immunofluorescence

Cells were plated on bovine fibronectin-coated (1:75 in PBS, Sigma-Aldrich F1141, CAT#F1141) or collagen-coated (1:100 in PBS, Advanced Biomatrix Cat#5005-B), 96-well, glass-bottom plates (Cellvis CAT# P96-1.5h-N). Cells were fixed in 4% PFA (in PBS, Fisher Scientific Cat#AA433689) for 15 min at room temperature (RT), washed 3x with PBS, permeabilized in 0.2% Triton X-100 for 15 min, and blocked in blocking solution (10% FBS, 5% BSA, 0.1% Triton X-100, 0.01% NaN3 in PBS) for 1h at RT. For conditions with EdU incorporation (10 μM in basal medium, 12-15 min at 37C, Cayman Chemical Cat#20518) prior to fixation, cells were washed 1x with PBS after the blocking step and treated with a click chemistry solution (2 mM CuSO4, 20 mg/mL sodium ascorbate, 3 μM AFDye picolyl azide (488, 568, or 647, Click Chemistry Tools Cat# 1276, 1292, or 1300 respectively) in TBS (50 mM Tris, 150 mM NaCl, pH 8.3) for 25 min. Following a 1x PBS wash, cells were incubated with primary antibody for 3h at room temperature or overnight at 4C (see also Table S2) in blocking solution, then washed 3x in PBS. Fluorophore-conjugated secondary antibodies (Thermo Fisher Scientific, goat anti-mouse, goat anti-rabbit, 1:1000 in blocking solution) and Hoechst 333342 (Thermo Fisher Scientific Cat#H3570, 1:10,000) were incubated on cells for 1h at RT, then washed 3x in PBS. Wells were imaged in 1x PBS or 1x N-acetylcysteine (NAC, 700 mM in ddH2O, pH 7.4, Sigma-Aldrich A7250) imaging buffer for iterative immunofluorescence protocol^32^ (4i).

For the 4i protocol, wells were then washed 1x in dH2O, treated with elution buffer (0.5M L-glycine (Sigma-Aldrich Cat#G2879), 3M urea (Sigma-Aldrich Cat#U5378), 3M guanidine hydrochloride (Sigma-Aldrich Cat# G3272), 70 mM TCEP-HCl (Goldbio Cat# TCEP50) in ddH2O, pH 2.5) for 20 min on an orbital shaker. After 1x PBS wash, elution was confirmed by fluorescence microscopy. Cells were then blocked with 4i blocking solution (1% BSA + 150 mM maleimide (Sigma-Aldrich Cat# 129585) in PBS) for 30 min, then incubated with the second round of primaries (in blocking solution) for at least 3h RT or overnight at 4C. Secondary staining, imaging in NAC, and subsequent elution and 4i blocking steps were repeated up to 5 times. Control wells with secondary antibody treatment alone were used to confirm proper elution of previous antibody rounds. Antibody list and dilutions used for immunofluorescence are listed below and in Table S2:

### Immunofluorescence Antibodies

The following primary antibodies were used exclusively for round 1 of immunofluorescence staining: rabbit anti-CDK6 (1:1000, Abcam Cat#ab124821, RRID: AB_10999714), rabbit anti-YAP (1:400, Cell Signaling Technology Cat# 14074, RRID:AB_2650491), mouse anti-p27 (1:1000, Cell Signaling Technology Cat#3698, RRID:AB_2077832), mouse anti-p21 (1:500, BD Bioscience Cat#556430, RRID:AB_396414), rabbit phospho-AKT(S473)(1:400, Cell Signaling Technology Cat#4060, RRID:AB_2315049), rabbit phospho-MAPK1/2(Thr202/Tyr204)(1:400, Cell Signaling Technology Cat#4370, RRID:AB_2315112), mouse phospho-S6(235/236)(1:800, Cell Signaling Technology Cat#62016, RRID:AB_2799618).

The following primary antibodies were used for any round of immunofluorescence: mouse anti-cyclin D1 (1:200, Santa Cruz Biotech Cat#sc-8396, RRID:AB_627344), mouse anti-FRA1 (1:200, Santa Cruz Biotech Cat#sc-28310, RRID: AB_ AB_627632), rabbit anti-Myc (1:800, Cell Signaling Technology Cat#5605, RRID:AB_1903938), mouse anti-Myc tag (1:8,000, Cell Signaling Technology Cat#2276, RRID:AB_331783), mouse anti-N-cadherin (1:200, Santa Cruz Biotech Cat#sc-393933, RRID: AB_2832921), rabbit anti-p27 (1:1600, Cell Signaling Technology Cat#3686, RRID:AB_2077850), rabbit anti-p21 (1:2500, Cell Signaling Technology Cat#2947, RRID:AB_823586), rabbit anti-phospho-Rb(Ser807/811)(1:2500, Cell Signaling Technology Cat#8516, RRID:AB_1117658), rabbit anti-phospho-S6(240/244)(1:2500, Cell Signaling Technology Cat#5364, RRID:AB_10694233), rabbit anti-Skp2 (1:800, Cell Signaling Technology Cat#2652, RRID:AB_11178941), mouse anti-YAP (1:200, Santa Cruz Biotech. Cat#sc-101199,RRID: AB_1131430). Antibodies were previously validated in RPE1^22^ and for 4i^81^, and in this study by drug treatment (including AKT, mTORC1/2, MEK, and LATS1/2 inhibitors).

The following secondaries were used in variable combination (488, 514 or 568, and 647), at 1:1000: goat anti-mouse IgG Alexa Fluor 488 (Thermo Fisher Scientific Cat#A11029, RRID:AB_2534088), goat anti-mouse IgG Alexa Fluor 514 (Thermo Fisher Scientific Cat#A31558, RRID:AB_1037789), goat anti-mouse IgG Alexa Fluor 568 (Thermo Fisher Scientific Cat#A-11031 RRID:AB_144696), goat anti-rabbit IgG Alexa Fluor 568 (Thermo Fisher Scientific Cat#A11036, RRID:AB_1056366), goat anti-rabbit IgG Alexa Fluor 647 (Thermo Fisher Scientific Cat#A21245, RRID:AB_23813).

### Microscopy

For automated epifluorescence microscopy, cells were imaged on a Ti2-E inverted microscope (Nikon) using penta-band (DAPI/CFP/YFP/TRITC/Cy5, Chroma: 89405) Sedat filter sets with an LED light source (Lumencor Spectra X or Spectra III), Hamamatsu ORCA-fusionBT or ORCA-Flash4.0 V3 sCMOS camera, and 20x objective (Nikon CFI Plan Apo Lambda, 0.75 NA) in a humidified chamber at 37 °C and 5% CO2. 25-30 tiled images were acquired per well in 16-bit mode with 2×2 binning. Fixed cells were imaged in 1x N-acetylcysteine imaging buffer. Fluorophores for secondary antibodies were previously optimized to minimize bleed-through^77^. For live-cell imaging, 9 sites were imaged per well and stitched into a single image to maximize the number of cells that could be tracked over time (with at least 3 well-replicates per experiment). Images were acquired at 12 min intervals of with the minimum laser intensity and light exposure to limit phototoxic effects and automated shading correction for each channel based on the autofluorescence of media containing 10% serum. For LATS1/2 and TEAD inhibitor experiments, asynchronously cycling cells were imaged in DMSO for 12h, then acutely treated with inhibitor and imaged for 24h to assess cell cycle entry and exit.

### RNA sequencing experimental conditions

WT and doxycycline-inducible YAP5SA cells were seeded at 38,000 cells per well of 96-well, optical-grade plates (Sigma-Aldrich Cat#CLS3904) in 1.0% FBS. After ∼2h to allow time for adherence to the tissue culture surface, cells were washed with 0.1% FBS then pre-arrested with CDK4/6 inhibitor (palbociclib, 1 uM, SelleckChem CAT#S1116) in either 1.0% FBS or 0.1% FBS for 28-36h prior to doxycycline (500 ng/mL, Sigma-Aldrich CAT#D9891) induction or DMSO control treatment for 6, 12, 24, or 36h in the presence of CDK4/6 inhibitor to maintain cells in G0 and prevent additional transcriptional variation due to cell cycle progression. Each experiment (repeated on three separate occasions) was performed in technical duplicate (with 6 replicate wells per technical duplicate), reserving one replicate for immunofluorescence staining to confirm quality of cell cycle arrest (by DNA content and EdU incorporation) and the other for harvesting RNA.

### RNA-seq and analysis

Prior to harvesting, 96-well plates were washed with PBS, aspirated fully and transferred immediately to −80C. RNA was later harvested from the plates by briefly thawing prior to using extraction by RNeasy Mini kit (Qiagen Cat# 74106) and QIAshredder (Qiagen Cat#79654), with pooling of 6 technical replicate wells/condition directly following the lysis step prior to following the manufacturer protocol, including the optional DNase treatment (Qiagen Cat# 79254). Samples were eluted in ultraPure ddH2O and assessed for quality using an Agilent Bioanalyzer TapeStation. Illumina Stranded mRNA library preparation with polyA capture and sequencing was performed on a NovaSeq6000 system (paired-end, 2 x 50) at the Weill Cornell Genome Resources Core Facility over two rounds to get sufficient read depth.

Raw reads were checked for quality using FASTQC then aligned to the human genome assembly (GRCh38) with HISAT2^82^ and assigned to genomic features with featureCounts^83^. Differential expression analysis was conducted with DESeq2^84^.

### Image quantifications and analysis

Tracking of cells for time-lapse experiments, illumination bias correction, nuclear segmentation, image cropping and alignment across iterative rounds of immunofluorescence, and analysis were performed with a custom MATLAB pipeline (previously published^77^ and deposited on Github: https://github.com/MeyerLab/image-analysis-ratnayeke-2021) incorporating local density measurements^22^. For illumination bias correction, images of background autofluorescence from immunofluorescence blocking solution in cell-free wells were taken for each experiment and used to flatten raw images. Cell nuclei were automatically segmented using expression of a fluorescently-tagged nuclear marker (H2B-miRFP670) or Hoechst staining, a Gaussian filter, and a curvature-based object splitting algorithm. A segmentation mask from the first round of imaging was used to align subsequent imaging rounds and crop the images for the overlapping regions. Image jitter between 4i rounds was typically less than 500 nm.

For nuclear-localized protein stains (EdU, cyclin D1, p27, phospho-Rb, c-Myc, Fra1, SKP2, CDK6, p21), global background subtraction was performed automatically by expanding a 7.8 μm ring from the nuclear mask and subtracting the 25^th^ percentile of pixel signal as estimated background. For protein stains with cytoplasmic and nuclear localization (YAP, phospho-S6, phospho-AKT, phospho-ERK, N-cadherin), median background signal was estimated during analysis by manual averaging of background pixels of representative cell-free regions and subtracting as a flat threshold. For quantification of immunofluorescence signals, DNA content was calculated as the total nuclear Hoechst intensity (median intensity x nuclear area); nuclear fluorescence signals were calculated as the median nuclear intensity; cytoplasmic fluorescence signals were calculated as the median cytoplasmic intensity (expanding the nuclear mask with a ring of inner radius 0.65 um and outer radius 3.23 um to minimize the chance of sampling cytoplasm of neighboring cells). To compare experimental replicates, fold-change in immunofluorescence staining was used to account for the variation in staining and real fluorescence unit values.

Cell gating for G1-phase cells was performed by distinguishing G0/G1 and G2 cell populations by 2N or 4N DNA context, then selecting cells below the EdU-incorporation threshold (see sample EdU histograms, Figure S1A) with 2N DNA. Otherwise, for calculating percent EdU-positive or percent Rb phosphorylated (all-cell gating), cells across cell cycle phases were analyzed together. Rb phosphorylation was defined by histogram analysis of the bifurcated phospho-Rb stain and calculating the fraction of cells above the threshold cutoff (see sample p-Rb histograms, Figure S1A).

For analysis of CDK2 activity at 3h following mitosis in cells with variable time spent in LATS1/2 or TEAD inhibitor (Figure 3K-L cell traces were first gated based on cells that were tracked for at least 20% of the total imaging time (36h) and tracked from the time of birth or mitosis (denoted as a daughter cell). CDK2 activity of the kinase translocation reporter was measured as a ratio of the median intensity of a cytoplasmic ring (0.65 – 3.25 μm expansion from the nuclear mask) over the median intensity of the nuclear signal. For calculating the fraction of cells with high CDK2 activity, asynchronously cycling cells were first binned into 3h windows by time of mitosis (to ensure enough cells analyzed per time window of mitosis) and then gated by time spent in the inhibitors by selecting cells that were born at variable times following the switch from DMSO to the LATS or TEAD inhibitor. The fraction of cells with CDK2 activity above a standardized threshold (0.8) was calculated 3h after mitosis as an early indicator of cells bifurcating into low or high activity^45^, but similar results were also seen if assessing CDK activity at 6h after mitosis (data not shown).

### Local cell density analysis

Local cell density analysis was adapted from Fan et al., 2021. Briefly, local density was measured by dividing imaging sites (25-30 tiled images/well) into a 3D histogram based on the centroid coordinates of segmented nuclei (10 x 10 grid, MATLAB hist3). The xy location coordinates of each cell centroid were rounded and the number of cells per block was automatically counted, then scaled back to match the original image size (giving numerical output rather than integer values)^22^. The local cell density value assigned for each cell was based on the scaled number of cells in its assigned local density block.

### Quantification corrections

While cells were counted prior to seeding, final cell plating density varied across experiments, and absolute proliferation levels varied per independent experiment. We therefore assessed both absolute (% EdU, % p-Rb) and fold-change proliferation metrics for assessing drug and condition effect sizes.

During the iterative steps of multiplexed immunofluorescence, some cells washed off during later rounds of imaging. As a proxy for null cells, cells that were segmented from the first round of imaging were gated out of the final analysis if their Hoechst value was less than 100 RFU for a subsequent imaging round.

For live-cell imaging experiments, some cells lost expression of the reporters of cell cycle activity and were therefore removed from analysis.

### Statistical Analysis

Statistical analyses were performed using Student’s one-sample or paired (MATLAB ttest) and two-sample (MATLAB ttest2) t tests. Linear regressions for data binned by local cell density were performed using fitlm (MATLAB). 95% confidence intervals and SEM are plotted as indicated for representative experiments and summary data.

## Supplemental information titles and legends

**Figure S1, related to Figure 1.**
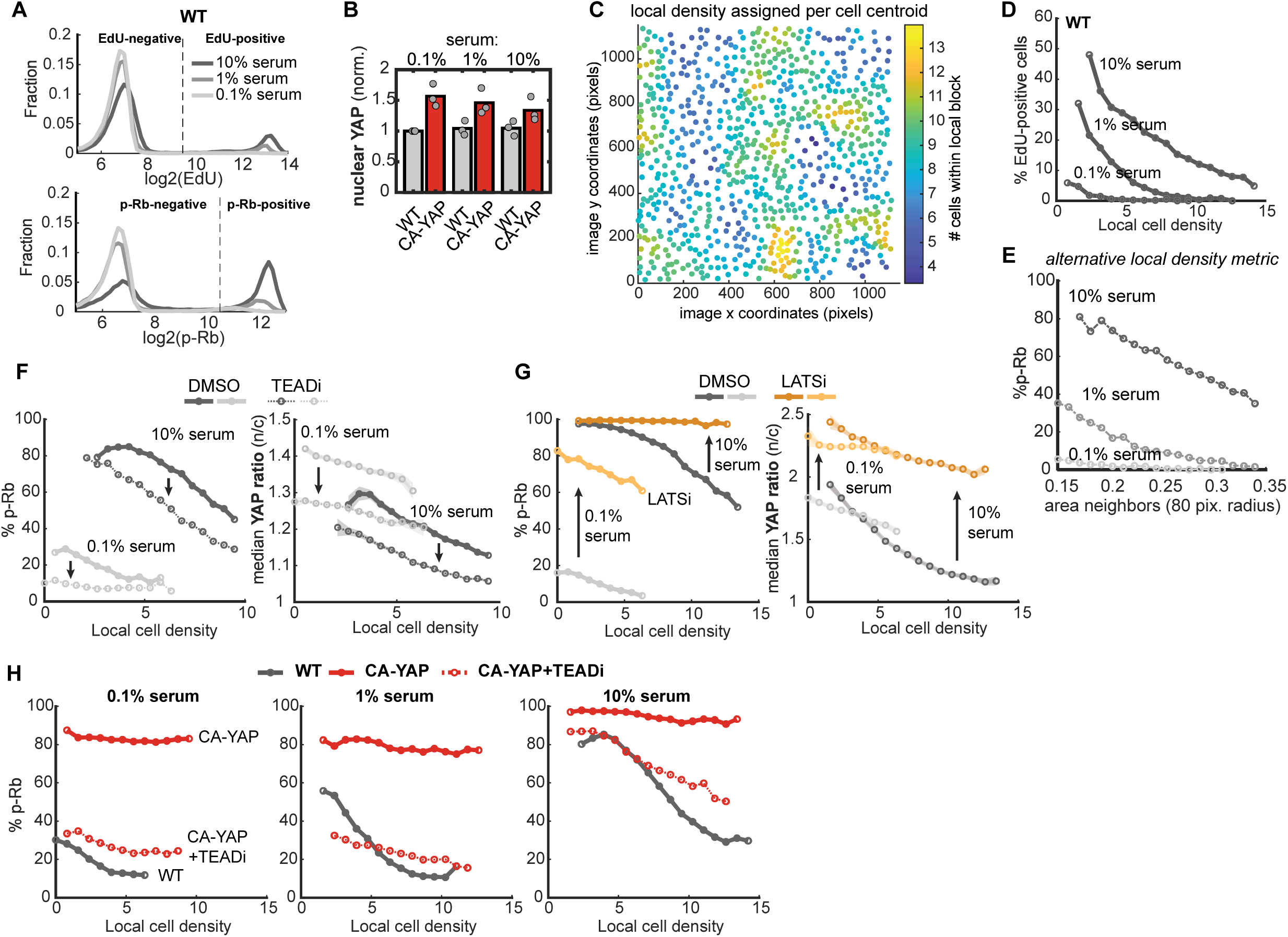
**A)** Histogram plots of binary distribution of EdU (<u>top</u>) and Rb phosphorylation (phospho-Rb 807/811, <u>bottom</u>) in WT RPE1 cells with 10%, 1%, or 0.1% serum. Dashed line indicates cutoff values used to designed whether cells are EdU-positive or have phosphorylated Rb. Data are representative of 3 independent experiments (same experiment as in Figure 1E). **B)** Mean nuclear YAP protein levels in serum-titrated WT and CA-YAP cells, normalized (norm.) to WT/0.1% serum condition for each experiment, n = 3 independent experiments. **C)** Representative local density estimation in an imaging site, where circles represent the location of the centroids of cell nuclei. Cell location coordinates were divided into a 10×10, equally-spaced grid (local density blocks). The z-scale color indicates the number of cell centroids within a block. Cells were assigned local densities based on the number of cells in their block, scaled by the image size to give a relative density unit. See also Methods. **D)** Percent EdU-positive cells per local density bin for WT cells in titrated serum levels. Data are representative of 3 independent experiments. **E)** Percent p-Rb binned by alternate local density metric (area of neighboring cell nuclear pixels within an 80-pixel (∼51.6 μm) radius) for WT cells in titrated serum, showing similar density dependence as the primary local density metric. **F-G)** Percent p-Rb (left) and median YAP ratio (nuclear/cytoplasmic, n/c, right) binned by local density for WT and CA-YAP cells in 0.1% and 10% serum. Cells were treated with DMSO and TEADi (1 μM GNE-7883, F, dashed lines) or LATSi (0.5 μM TDI-011536, G, orange lines) for 24h. Shaded error are 95% confidence intervals, n > 15,000 cells/condition. Arrows highlight drug-mediated changes for matched serum conditions. **H)** Percent p-Rb per local density bin for WT (grey) and CA-YAP cells (red) treated with DMSO or TEADi (0.5 μM, dashed line) in titrated serum. All binned and histogram data are from one representative replicate of 3 independent experiments, n >100 cells/bin and n >15,000 cells/condition).

**Figure S2, related to Figure 2.**
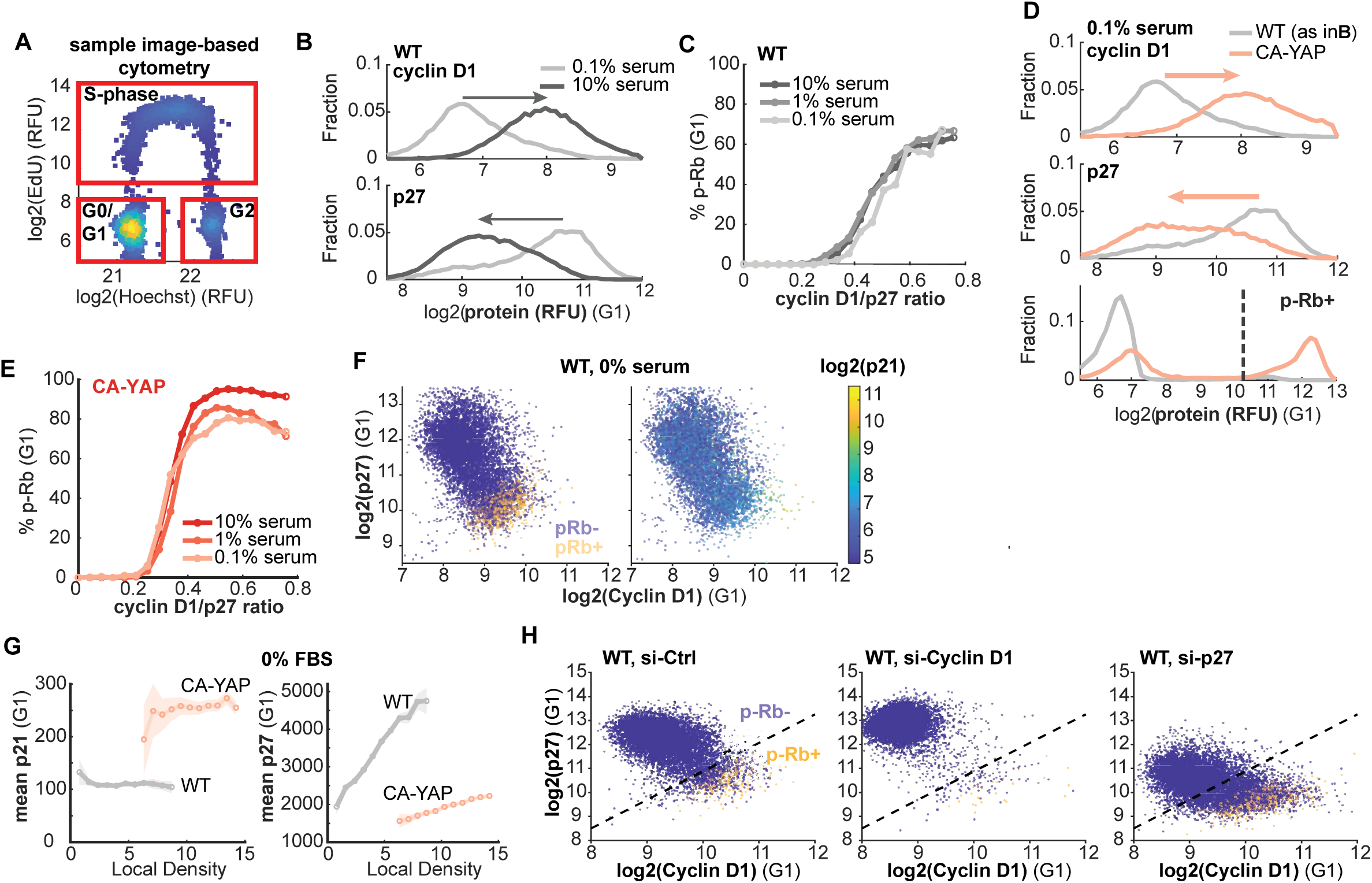
**A)** Representative density scatterplot of DNA content (Hoechst staining, real fluorescence units (RFU)) and EdU-incorporation for G0/G1 gating of cells. G1 population cells were defined as 2N DNA/EdU-negative, S-cells as EdU-positive, and G2 as 4N DNA/EdU-negative. All-cell gating reflects cells within the outer red boundary (n = 8,000 random cells). **B)** Histogram distribution of cyclin D1 protein (<u>top</u>) and p27 protein (<u>bottom</u>) stain in WT cells in 0.1% or 10% FBS. **C)** Percent p-Rb binned by cyclin D1/p27 ratio for serum titration in WT cells (n > 15,000 cells/condition). **D)** Histogram distribution of cyclin D1 (<u>top</u>), p27 (<u>middle</u>), and p-Rb (<u>bottom</u>) stain comparing CA-YAP cells with WT in 0.1% FBS (same WT data as S1A), n > 20,000 cells/condition. Dashed line indicates p-Rb cutoff. **E)** Percent p-Rb binned by cyclin D1/p27 ratio for serum titration in CA-YAP cells (n > 15,000 cells/condition). **F)** Representative density scatterplot of cyclin D1 stain against p27 stain, colored by binary p-Rb status (left) or p21 stain (right, parula scale) in starved WT cells. **G)** Mean p21 (left) and p27 (right) protein levels (RFU) binned by local density for starved WT (grey), and CA-YAP (red) cells (n > 20,000 cells/condition). Shaded error are 95% confidence intervals. **H)** Representative density scatterplots of cyclin D1 and p27 protein levels (RFU) colored by p-Rb status for serum-starved WT cells transfected with siRNA against control (si-Ctrl, left), cyclin D1 (si-Cyclin D1, middle), or p27 (si-p27, right). Dashed, diagonal line visually approximates where p-Rb bifurcates in CA-YAP conditions (matching Figure 2H). All data representative of 2 independent experiments and gated for G1-phase. All single-cell scatterplots show n = 8,000 cells.

**Figure S3, related to Figure 3.**
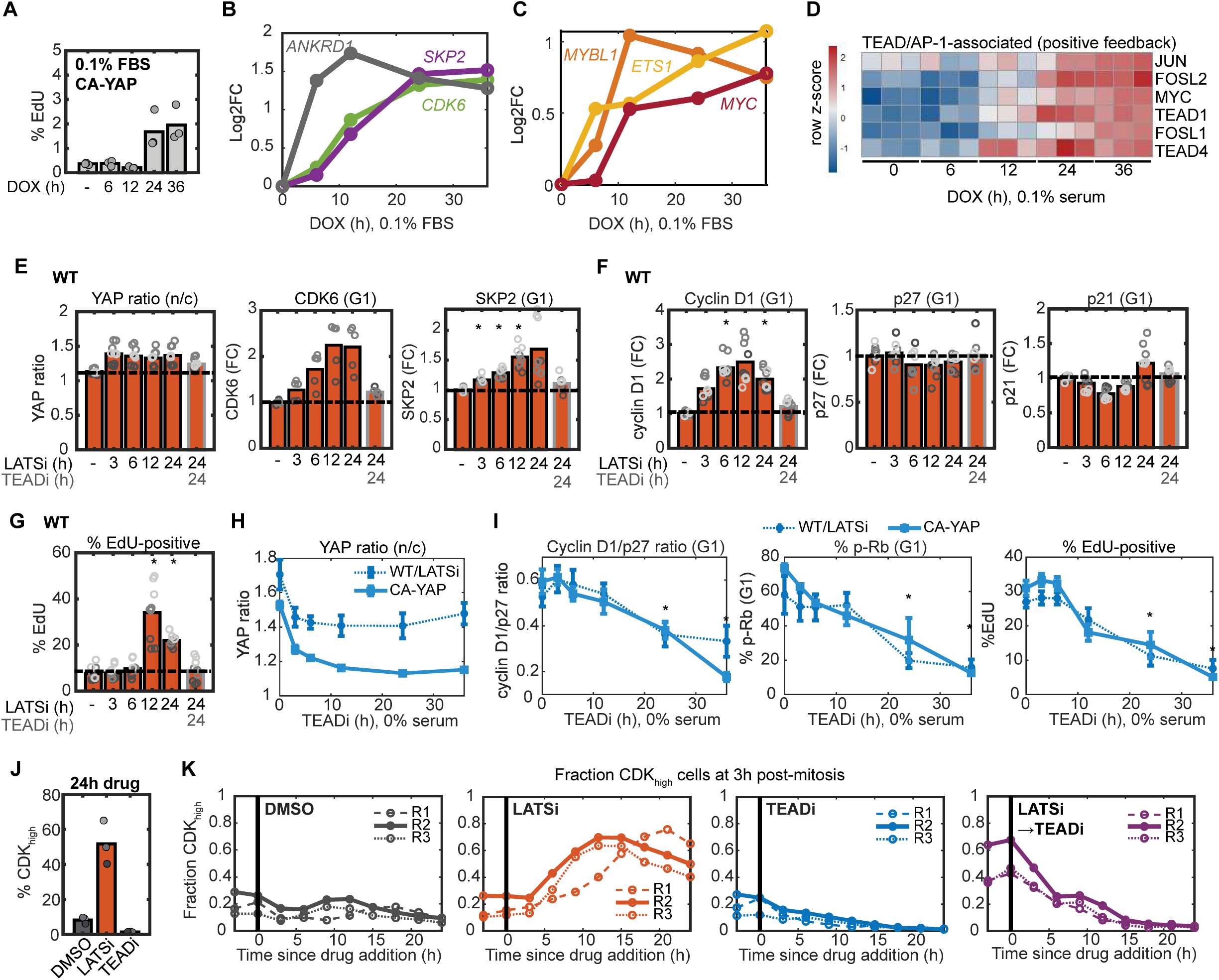
**A)** Mean percent EdU for matched replicates of Figure 3A reserved for immunofluorescence staining for CA-YAP cells in 0.1% FBS and arrested with palbociclib (1μM) during treatment with DMSO (-) or doxycycline (DOX) for 0-36h. N = 3 independent experiments. **B-C)** Kinetics of sample YAP target gene induction. Log2 fold-change (Log2FC) induction of ANKRD1 (canonical YAP target gene, grey) compared with *CDK6* (green) and *SKP2* (purple) (B) and transcription factor targets *MYC* (red) compared with *MYBL1* (orange) and *ETS1* (yellow) at times since overexpression (C). Adjusted p-values: *CDK6*, 0.0451 (6h), 4.22 x10^-22^ (12h), 3.08 x10^-52^ (24h), 3.26 x10^-58^ (36h); *SKP2*, 3.78 x10^-13^ (12h), 3.49 x10^-65^ (24h), 1.86 x10^-70^ (36h); *MYC*, 2.18×10^-6^(12h), 2.39×10^-8^ (24h), 1.50×10^-13^ (36h); *MYBL1*, 7.77×10^-13^ (12h), 2.76×10^-10^ (24h), 3.48×10^-7^ (36h); *ETS1*, 1.85×10^-13^ (6h), 4.55 x10^-16^(12h), 6.24×10^-37^(24h), 1.23×10^-57^(36h). **D)** Heatmap of TEAD/AP-1-associated positive feedback genes in DOX-treated, CA-YAP cells. Color scale indicates row z-score. **E-G)** Mean YAP ratio (E, left) and fold-change in protein levels of CDK6 (E, middle), SKP2 (E, right), cyclin D1 (F, left), p27 (F, middle), p21 (F, right), and percent EdU (G) for serum starved, WT cells treated with LATSi (0.5 μM TDI-011536) for 0-24h or co-inhibited with TEADi (grey outline, 1 μM GNE-7883). N = 2 (CDK6) or 3 independent experiments (all other proteins). Student’s one-sample t-test: SKP2, p = 0.0159 (3h), 0.0178 (6h), 0.0468 (12h); cyclin D1, p = 0.0168 (6h), 0.0271 (24h); EdU, p = 0.0285 (12h), 6.65×10^-4^ (24h). **H-I)** Mean YAP ratio (H), cyclin D1/p27 ratio (I, left), percent p-Rb (I, middle) and EdU-positive (I, right) cells following TEADi treatment of starved WT cells treated with LATSi (dashed line) or CA-YAP (solid line, same data as Figure 3I). Mean ± SEM (n = 3 independent experiments). Student’s two-sample t-test, CA-YAP: cyclin D1/p27 ratio, p = 0.0354 (24h, 36h); %p-Rb, p = 0.0377 (24h), 0.0328 (36h); EdU, 0.0038 (24h), 0.0012 (36h). **J)** Mean percentage CDK_high_ cells at 3h post-mitosis for cells reentering the cell cycle 24h after DMSO, LATSi, or TEADi treatment. N = 3 independent experiments. **K)** Fraction of CDK_high_ cells at 3h post-mitosis (mid-G1 phase) relative to drug treatment. Cells were aligned by mitosis and binned based on time spent in control (DMSO), LATSi, or TEADi (n > 100 cells/bin, n > 5,000 cells/condition). For the LATSi to TEADi drug change (right), cells were treated with LATSi for 12h prior to washout and treatment with TEADi. Each line represents an independent experiment (n = 3), where matched line styles indicate data from the same experiment. All fixed-cell data are G1-gated. Colored rings indicate replicate wells within matched experimental replicates.

**Figure S4, related to Figure 4.**
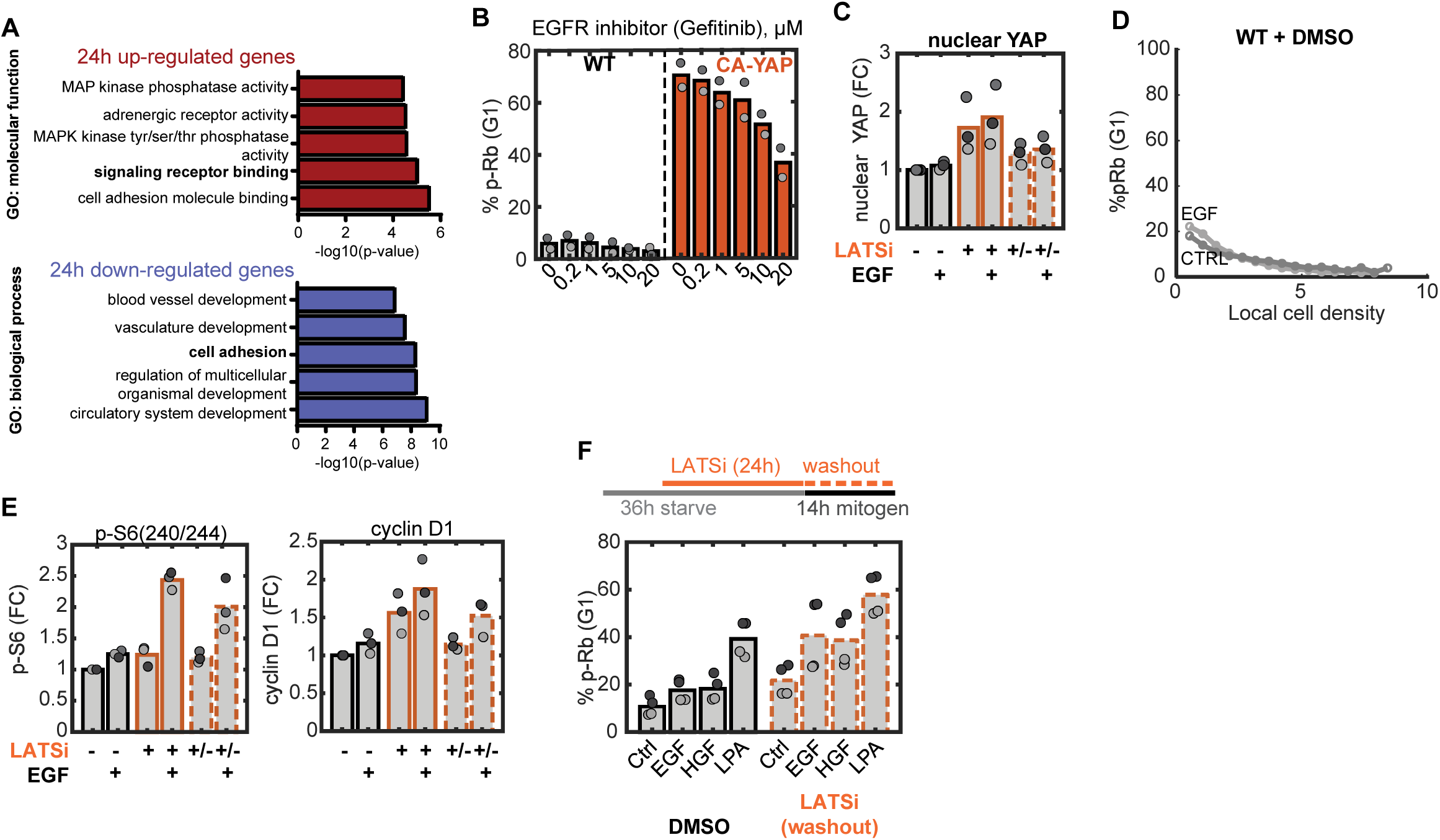
**A)** <u>Top</u>: GO terms enriched for significantly up-regulated genes after 24-hour YAP induction (q < 0.05, Log2FC > 1.0) relative to control (0.1 % FBS) (Figure 5A). <u>Bottom</u>: GO process terms enriched for significantly down-regulated genes (q < 0.05, Log2FC < −1.0). **B)** Mean percent p-Rb for serum-starved WT and CA-YAP cells treated with gefinitib (EGFR inhibitor) at indicated doses (μM) for 24h (n = 2 independent experiments). **C)** Mean fold-change in nuclear YAP levels for WT cells treated with DMSO or LATSi ± washout during EGF treatment (0.5 μM TDI-011536, 25 ng/mL EGF) as in Figure 4G (n = 3 independent experiments). **D)** Percent p-Rb binned by local density for serum-starved WT cells pre-treated with DMSO prior to 14h EGF (25 ng/mL) or control (CTRL) treatment, as in Figure 4G. Data are representative of 3 independent experiments (n > 25,000 cells/condition). **E)** Mean fold-change in cytoplasmic p-S6 (240/244, <u>left</u>) and cyclin D1 (<u>right</u>) for cells treated as in Figure 4G (n = 3 independent experiments, where matched colors indicate values from same experiment). **F)** Mean percent p-Rb for serum-starved WT cells pre-treated with DMSO or LATSi (5 μM, TRULI) for 24h prior to washout and 14h mitogen treatment (25 ng/mL EGF and HGF, 1 μM LPA). N = 2 independent experiments, where matched colors indicate well-replicates/experiment. All single-cell data are G1-gated cells.

**Figure S5, related to Figure 5.**
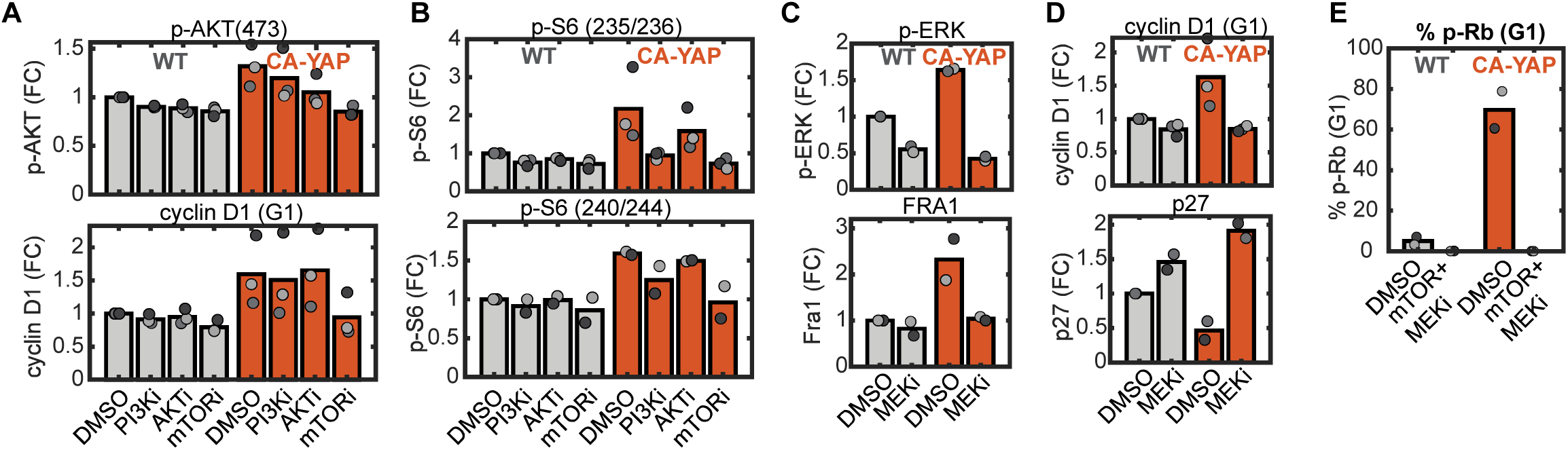
**A-B**) Fold-change in protein levels of p-AKT(S473) and cyclin D1 (A), and p-S6(B, <u>top</u>: 235/236; <u>bottom</u>: 240/244) for serum-starved WT and CA-YAP cells treated with PI3Ki, AKTi, or mTORi, as in Figure 5E. N = 2 (p-AKT, p-S6-240/244) or n = 3 (cyclin D1, p-S6-235/236) independent experiments, where matched colors indicate values from the same experiment. **C-D**) Mean fold-change in protein levels of p-ERK and Fra1 (C), cyclin D1 and p27 (D) for serum-starved WT and CA-YAP cells treated with MEKi as in Figure 5F. N = 2 (p-ERK, Fra1, p27) or n = 3 (cyclin D1) independent experiments, matched colors indicate values from the same experiment. **E)** Mean percent p-Rb in serum-starved WT and CA-YAP cells treated with DMSO or combined mTOR+MEKi (100 nM Torin2, 100 nM PD0325901). N = 2 independent experiments, matched colors indicate values from the same experiment. All data are from G1-gated cells.

**Figure S6, related to Figure 6.**
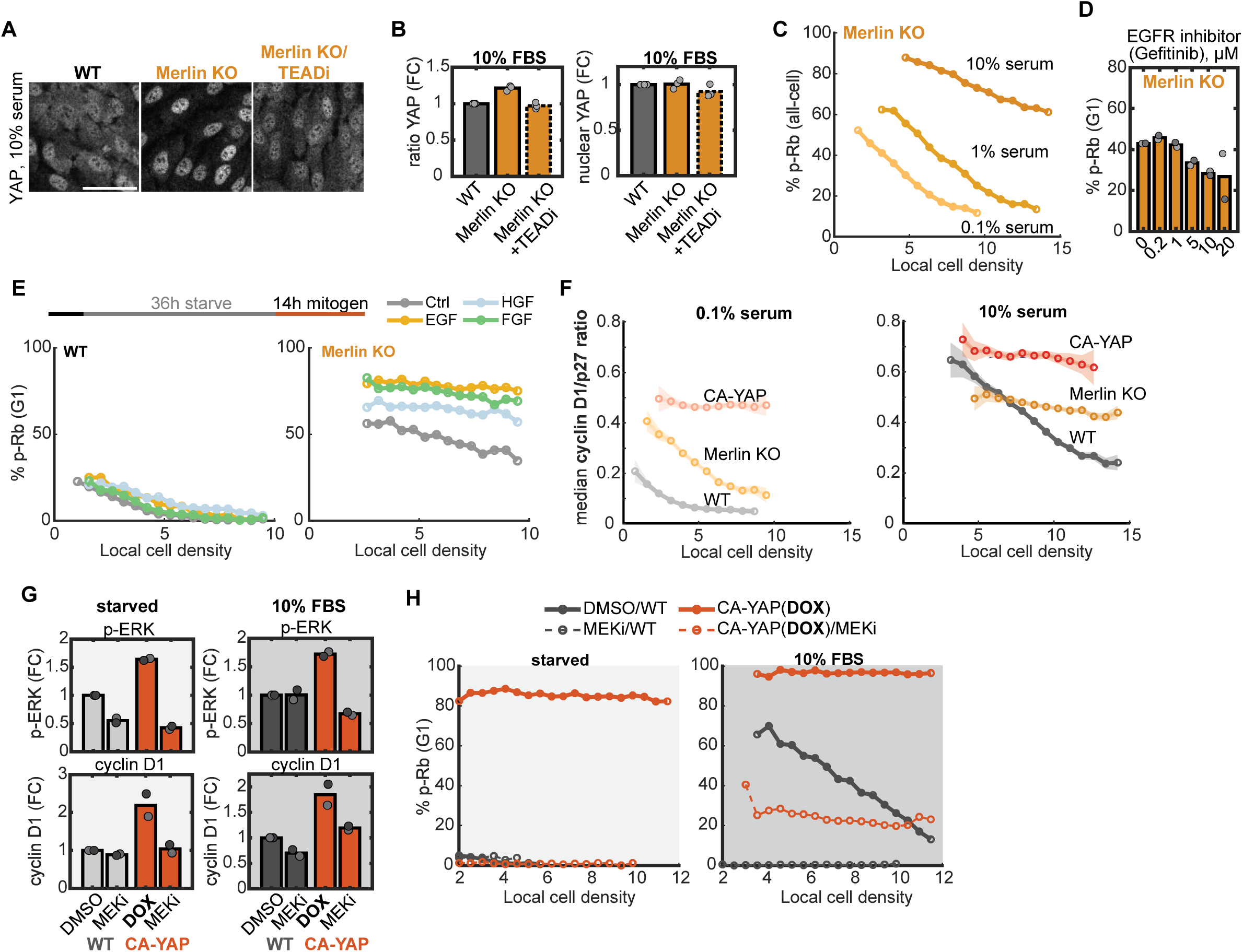
**A)** Representative immunofluorescence images of YAP staining in WT, Merlin KO, and TEADi-treated Merlin KO (0.5 μM GNE-7883) in 10% serum. Scale bar = 50 μm. **B)** Mean fold-change in YAP ratio and nuclear YAP levels in Merlin KO treated with DMSO or TEADi relative to WT cells, as in (A). N = 3 independent experiments. **C)** Percent p-Rb per local cell density bin for Merlin KO cells maintained in titrated serum levels (gating for all cells, as in Figure 1C). N > 20,000 cells/condition, data are representative of 3 independent experiments. **D)** Mean percent p-Rb for serum-starved Merlin KO cells treated with gefitinib (EGFR inhibitor) at indicated doses (μM) for 24h. N = 2 independent experiments. **E)** <u>Top</u>: starvation-release protocol for WT or Merlin KO cells with control (CTRL), EGF, HGF, or FGF (all 25 ng/mL). <u>Bottom</u>: percent p-Rb comparing WT (left) and Merlin KO (right) treated with CTRL or mitogens for 14h. N > 12,000 cells/condition, data are representative of 2 independent experiments. **F)** Median cyclin D1/p27 ratio binned by local cell density for WT (grey), Merlin KO (yellow), and CA-YAP cells (red) in 0.1% (left) or 10% serum (right). Shaded error are 95% confidence intervals, data are representative of 2 independent experiments. **G)** Mean fold-change in p-ERK (<u>top</u>) and cyclin D1 (<u>bottom</u>) for WT and DOX-induced CA-YAP cells treated as in Figure 5F in starvation (left) and 10% serum conditions (right). N = 2 independent experiments, where matched colors indicate values from the same experiment. **H)** Percent p-Rb binned by local density for WT and DOX-induced CA-YAP cells starved (left) or in 10% serum (right) treated with DMSO or MEKi as in (G). Starved: n > 5,000 cells (WT), n > 20,000 cells/condition (CA-YAP); 10% serum: n > 10,000 cells (WT), n > 20,000 cells/condition. Data are representative of 2 independent experiments. All binned data are G1-gated unless stated otherwise, with n > 100 cells/bin.

