## SupplementalTablesS1-S2 for "Transient proliferation by reversible YAP and mitogen-control of the cyclin D1/p27 ratio"

**Table S1**

| Oligo | Sequence (5’ 🡪3’) |  | Application |
| --- | --- | --- | --- |
| Merlin KO sgRNA (cloned into px459) | Forward: CACCGCACAGTGGCCTGGCTCAAAA  Reverse: AAACTTTTGAGCCAGGCCACTGTGC |  | Merlin KO generation |
| Pooled siRNA control | 1: GCCCUUCUCUGCAGUCAAG  2: UGGUUUACAUGUUGUGUGA  3: UGGUUUACAUGUUUUCUGA  4: UGGUUUACAUGUUUUCCUA | D-001810-10-05 | Transient transfection |
| Pooled siRNA CCND1 | 1: ACAACUUCCUGUCCUACUA  2: GUUCGUGGCCUCUAAGAUG  3: GCAUGUAGUCACUUUAUAA  4: GCGUGUAGCUAUGGAAGUU | L-003210-00-0005 | Transient transfection |
| Pooled siRNA CDKN1B (p27) | 1: CAGCTCGCTAGGAGCCGGGG  2: GGTCCAGGTCCCGGCTTCCC  3: GCGACCAGCCAATCTCCCGG  4: CGCTCGGGAACGAGGGGAGG | L-040178-00-0005 | Transient transfection |
| Pooled sgRNA non-targeting control | 1: GATACGTCGGTACCGGACCG  2: GTAACGCGAACTACGCGGGT  3: GTCGACGTTATTGCCGGTCG  4: GGAAACCTACGTCGACGAAT | U-009500-10-05 | Transient transfection |
| Pooled sgRNA CCND1 | 1: CCAAACGCCGGGAGCAGCGA  2: GAGCCCAAAAGCCATCCCTG  3: ATCAAAGCCCGGCAGAGAAT  4: GCCTGGGGAGACCACGAGAA | P-003210-01-0005 | Transient transfection |
| Pooled sgRNA CDKN1B (p27) | 1: CAGCTCGCTAGGAGCCGGGG  2: GGTCCAGGTCCCGGCTTCCC  3: GCGACCAGCCAATCTCCCGG  4: CGCTCGGGAACGAGGGGAGG | P-003472-01-0005 | Transient transfection |

**Table S2**

| Antibody | Vendor, Catalog # | Dilution | Application |
| --- | --- | --- | --- |
| Goat anti-mouse polyclonal secondary antibody (IRDye 800CW) | LI-COR Biosciences, CAT#925-32210 | 1:10,000 | Western blotting |
| Goat anti-mouse polyclonal secondary antibody (IRDye 680LT) | LI-COR Biosciences, CAT#925-68020 | 1:10,000 | Western blotting |
| Goat anti-rabbit polyclonal secondary antibody (IRDye 680LT) | LI-COR Biosciences, CAT#925-68021 | 1:10,000 | Western blotting |
| Goat anti-mouse polyclonal secondary antibody (IRDye 800CW) | LI-COR Biosciences, CAT#925-32211 | 1:10,000 | Western blotting |
| Goat anti-rabbit, HRP-linked secondary antibody | Cell Signaling Technology, CAT#7074 | 1:5,000 | Western blotting |
| Goat anti-mouse, HRP-linked secondary antibody | Cell Signaling Technology, CAT#7076 | 1:5,000 | Western blotting |
| Goat anti-mouse secondary antibody, Alexa Fluor 488 | Thermo Scientific,  CAT#A-11029 | 1:1,000 | Immunofluorescence |
| Goat anti-mouse secondary antibody, Alexa Fluor 514 | Thermo Scientific,  CAT#A-31555 | 1:1,000 | Immunofluorescence |
| Goat anti-mouse secondary antibody, Alexa Fluor 568 | Thermo Scientific,  CAT#A-11004 | 1:1,000 | Immunofluorescence |
| Goat anti-rabbit secondary antibody, Alexa Fluor 488 | Thermo Scientific,  CAT#A-11008 | 1:1,000 | Immunofluorescence |
| Goat anti-rabbit secondary antibody, Alexa Fluor 568 | Thermo Scientific,  CAT#A-11011 | 1:1,000 | Immunofluorescence |
| Goat anti-rabbit secondary antibody, Alexa Fluor 647 | Thermo Scientific,  CAT#A-21245 | 1:1,000 | Immunofluorescence |
| ANKRD1, rabbit polyclonal | Abcam, CAT#ab272894 | 1:2,000 | Western blotting |
| AXL (C89E7, rabbit monoclonal) | Cell Signaling Technology,  CAT#86616 | 1:1000 | Western blotting |
| CDK6, rabbit monoclonal | Abcam  CAT#ab124821 | 1:1000 | Immunofluorescence |
| c-myc (D84C12, rabbit monoclonal) | Cell Signaling Technology, CAT#5605 | 1:800 | Immunofluorescence |
| Cyclin D1 (A-12, mouse monoclonal) | Santa Cruz Biotechnology,  CAT#sc-8396 | 1:200 | Immunofluorescence |
| EGFR (D38B1, XP rabbit monoclonal) | CAT#4267 | 1:1,000 | Western blotting |
| Fra-1 (C-12, mouse monoclonal) | Santa Cruz Biotechnology  CAT#sc-28310 | 1:200 | Immunofluorescence |
| GAPDH  Rabbit monoclonal | Cell Signaling Technology, Cat#5174 | 1:5,000 | Western blotting |
| Myc epitope, mouse monoclonal | Cell Signaling Technology, Cat#2276 | 1:8,000 | Immunofluorescence |
| N-cadherin (H-2, mouse monoclonal) | Santa Cruz Biotechnology  CAT#sc-393933 | 1:300 | Immunofluorescence |
| P21 Waf1/Cip1 (SX118, mouse monoclonal) | BD Biosciences   CAT#556430 | 1:500 (IF, round 1 only) | Immunofluorescence |
| P21 Waf1/Cip1 (12D1, rabbit monoclonal) | Cell Signaling Technology,  CAT#2947 | 1:2500 (IF, any round) | Immunofluorescence |
| P27 Kip1 (SX53G8.5, mouse monoclonal) | Cell Signaling Technology,  CAT#3698 | 1:1000 (IF, WB) | Immunofluorescence |
| P27 Kip1 (D69C12, XP rabbit monoclonal) | Cell Signaling Technology,  CAT#3686 | 1:1600 | Immunofluorescence |
| Phospho-Akt (Ser473, D9E, XP rabbit monoclonal) | Cell Signaling Technology, CAT#4060 | 1:200 (IF) | Immunofluorescence, western blotting |
| Phospho-Erk1/2 (P44/42 MAPK Thr202/Tyr204) | Cell Signaling Technology,  CAT#4370 | 1:400 (IF)  1:1000 (WB) | Immunofluorescence, western blotting |
| Phospho-Rb (Ser807/811, D10B12, XP rabbit monoclonal) | Cell Signaling Technology, CAT#8516 | 1:2500 (IF)  1:1000 (WB) | Immunofluorescence, western blotting |
| Phospho S6 (Ser235/236, E2R10, mouse monoclonal) | Cell Signaling Technology,  CAT#62016 | 1:800 | Immunofluorescence |
| Phospho S6 (Ser240/244, D68F8, XP rabbit monoclonal) | Cell Signaling Technology, CAT#5364 | 1:2500 (IF) | Immunofluorescence |
| Skp2 (D3G5, XP rabbit monoclonal) | Cell Signaling Technology, CAT#2652 | 1:800 (IF) | Immunofluorescence |
| Vinculin (7F9, mouse monoclonal) | Thermo Scientific,  CAT#14-97777-80 | 1:1000 | Western blotting |
| YAP (63.7, mouse monoclonal) | Santa Cruz Biotechnology,  CAT#sc-101199 | 1:200 (IF, any round) | Immunofluorescence |
| YAP (D8H1X, XP rabbit monoclonal) | Cell Signaling Technology,  CAT#14074 | 1:400 (IF, round 1 only) | Immunofluorescence |
